# A trimeric USP11/USP7/TCEAL1 complex stabilizes RNAPII during early transcription to sustain oncogenic gene expression

**DOI:** 10.1101/2024.07.29.605622

**Authors:** Markus Dehmer, Peter Gallant, Steffi Herold, Giacomo Cossa, Francesca Conte, Jan Koster, Florian Sauer, Christina Schülein-Völk, Carsten P. Ade, Raphael Vidal, Caroline Kisker, Rogier Versteeg, Petra Beli, Seychelle M. Vos, Martin Eilers, Gabriele Büchel

## Abstract

During early transcription, RNA polymerase II (RNAPII) undergoes a series of structural transitions controlled by cyclin-dependent kinases. Whether protein ubiquitylation and proteasomal degradation affect the fate of RNAPII close to promoters is less well understood. Here we show that the deubiquitylating enzyme USP11 and its heterodimeric partner USP7 form a trimeric complex with TCEAL1, a member of the poorly understood TCEAL (TCEA/TFIIS-like) protein family. TCEAL1 shares sequence homology with the RNAPII interaction domain of the TCEA/TFIIS elongation factor, which controls the fate of backtracked RNAPII. TCEAL1 stabilizes complexes of USP11 with USP7 and with RNAPII. TCEAL1 is recruited to core promoters when transcription elongation is blocked and globally enhances the chromatin association of RNAPII during early transcription. Mechanistically, the USP11/USP7/TCEAL1 complex competes with TFIIS for binding to core promoters and protects RPB8, an essential subunit of RNAPII, from degradation, likely preventing excessive TFIIS-mediated transcript cleavage and RNAPII disassembly. In neuroblastoma and other tumors, TCEAL1-dependent genes define a TGF beta-dependent gene expression program that is characteristic for mesenchymal and invasive tumor cell types, suggesting that the USP11/USP7/TCEAL1 trimer stabilizes RNAPII during early transcription to support a critical oncogenic gene expression program (190 words).

## Introduction

In eukaryotic cells the synthesis of mRNAs is regulated by multiple mechanisms that control the function of RNA polymerase II (RNAPII) during the initial steps of transcription (Cramer 2019). Reflecting this complexity, RNAPII adopts a series of distinct structural states as it progresses from initial promoter binding to productive elongation (Vos et al. 2018). The transitions between these states involve changes in the phosphorylation of the carboxy-terminal domain (CTD) of RNAPII, which acts as a platform for the recruitment of factors that enable specific steps in RNAPII function (Schier and Taatjes 2020). One such transition is the release of RNAPII, which pauses after transcribing the first 20-60 nucleotides, into productive elongation (Core and Adelman 2019). Several specific factors, including NELF, DSIF, TFIIS and PAF1c, control pause release and transcription elongation. Among them, transcription factor II S (TFIIS, encoded by the *TCEA1*, *TCEA2* and *TCEA3* genes) has a specific role since it acts on RNAPII, which has backtracked on the DNA template (Noe Gonzalez et al. 2021). The interaction of TFIIS with RNAPII stimulates an endonucleolytic cleavage of the nascent transcript by RNAPII, which generates a new 3’-OH end and thereby allows transcription elongation to resume (Wind and Reines 2000). TFIIS can also promote the progressive degradation of nascent transcripts by RNAPII (Zatreanu et al. 2019). Furthermore the TFIIS interaction surface on RNAPII is also used by the repair factor UVSSA (Kokic et al. 2021) and is blocked by NELF in the paused state (Su and Vos 2024). Collectively, these observations argue that the interaction of TFIIS with RNAPII must be regulated *in vivo* to be productive.

Ubiquitylation of RNAPII and its subsequent proteasomal degradation play central roles in coordinating transcription with DNA repair and RNA metabolism, since they remove polymerase molecules that have encountered transcription-blocking lesions from chromatin, allowing for transcription-coupled DNA repair (Nakazawa et al. 2020; Tufegdzic Vidakovic et al. 2020). Several ubiquitin ligases, including NEDD4 (Anindya et al. 2007; Sun et al. 2013), the elongin/cullin-5 complex (CRL5^elongin^) (Yasukawa et al. 2008), and BRCA1 (Wu et al. 2007), can ubiquitylate RNAPII in response to different forms of DNA damage. The BRCA1, RNF168 and FANCI-FANCD2 ubiquitin ligases promote the removal of DNA-RNA hybrids (R-loops) that form between the nascent RNA and the coding strand of DNA when mRNA splicing and processing are impaired (Herold et al. 2019; Patel et al. 2021; Olazabal-Herrero et al. 2024). At the same time, multiple ubiquitin ligases degrade gene-specific or general transcription factors at promoters, arguing that ubiquitin ligase function needs to be restrained for some transcriptional programs and that this control may achieve specificity in gene regulation (Agricola et al. 2011; Bacon et al. 2020; Endres et al. 2021; Solvie et al. 2022).

Previous work has shown that two deubiquitylating enzymes, USP11 and USP7, associate with each other and are involved in multiple processes related to transcription and genomic stability (Sowa et al. 2009; Maertens et al. 2010). We therefore analyzed the USP11 interactome and found that USP11 interacts with TCEAL1 and TCEAL4, two members of the TCEA-like family of proteins (Yeh and Shatkin 1994; Pillutla et al. 1999). TCEAL1 shares a small region of sequence homology with the RNAPII-interaction domain of TFIIS (hereafter: <u>T</u>FIIS-<u>h</u>omology <u>s</u>equence, THS), but lacks an effector domain that enables TFIIS to alter RNAPII function, suggesting that it may act in a dominant-negative manner to buffer TFIIS function (Pillutla et al. 1999). During normal development, loss-of-function mutations in TCEAL1 cause an early-onset neurological disease trait consisting of hypotonia, neurobehavioral abnormalities, and dysmorphic facial features (Hijazi et al. 2022; Albuainain et al. 2024). The molecular mechanism of TCEAL1 action as well as potential roles of TCEAL1 in tumor biology remain largely unknown. Here we show that TCEAL1 and USP11 stabilize RNAPII on chromatin, that TCEAL1 competes with TFIIS for binding to RNAPII and that it is required to maintain RNAPII in a functionally active state to sustain a mesenchymal and TGF-beta dependent gene expression program that is conserved among many tumors.

## Results

### USP11 interacts with multiple proteins involved in basal transcription

To understand the function of USP11, we analyzed the interactome of USP11. We stably expressed HA-tagged USP11 at either the amino- or carboxy-terminus in human SH-EP neuroblastoma cells and performed mass spectrometry on anti-HA immunoprecipitates, using cells that do not express HA-tagged protein as negative control (Figure 1a,b and Supplemental Figure S1a,b and Supplemental Table 1). The experiments identified 342 proteins that were enriched in anti-HA immunoprecipitates from both cells expressing NT- or CT-tagged USP11 compared to immunoprecipitates from control cells (Supplemental Figure S1c). USP11 forms a heterodimer with USP7 (Sowa et al. 2009; Maertens et al. 2010), which is observed in both the mass spectrometry and the co-immunoprecipitation experiments (Figure 1a-d and Supplemental Figure S1d). Additionally, functional annotation showed that USP11 associates with multiple proteins involved in transcription elongation and the ubiquitin-proteasome system (Figure 1a,b and Supplemental Figure 1d). Unbiased gene ontology and pathway analyses confirmed that proteins in both functional categories were significantly enriched in the interactome (Supplemental Figure 1e). For example, two subunits of RNAPII (RPB1 and RPB2) as well as the elongation factor SPT6 (*SUPT6H*) (Narain et al. 2021) and several members of the Cullin family of ubiquitin ligases were present in the USP11 interactome (Figure 1a,b and Supplemental Figure 1d). Furthermore, we identified two proteins of unknown function, TCEAL1 and TCEAL4, as interactors of USP11 (Figure 1a, b). Both are members of the transcription factor elongating A-like family, which comprises nine nuclear phosphoproteins that are called TCEA-like proteins since some members, including TCEAL1 and TCEAL4, share homology to TFIIS (Yeh and Shatkin 1994; Pillutla et al. 1999).

**Figure 1:**
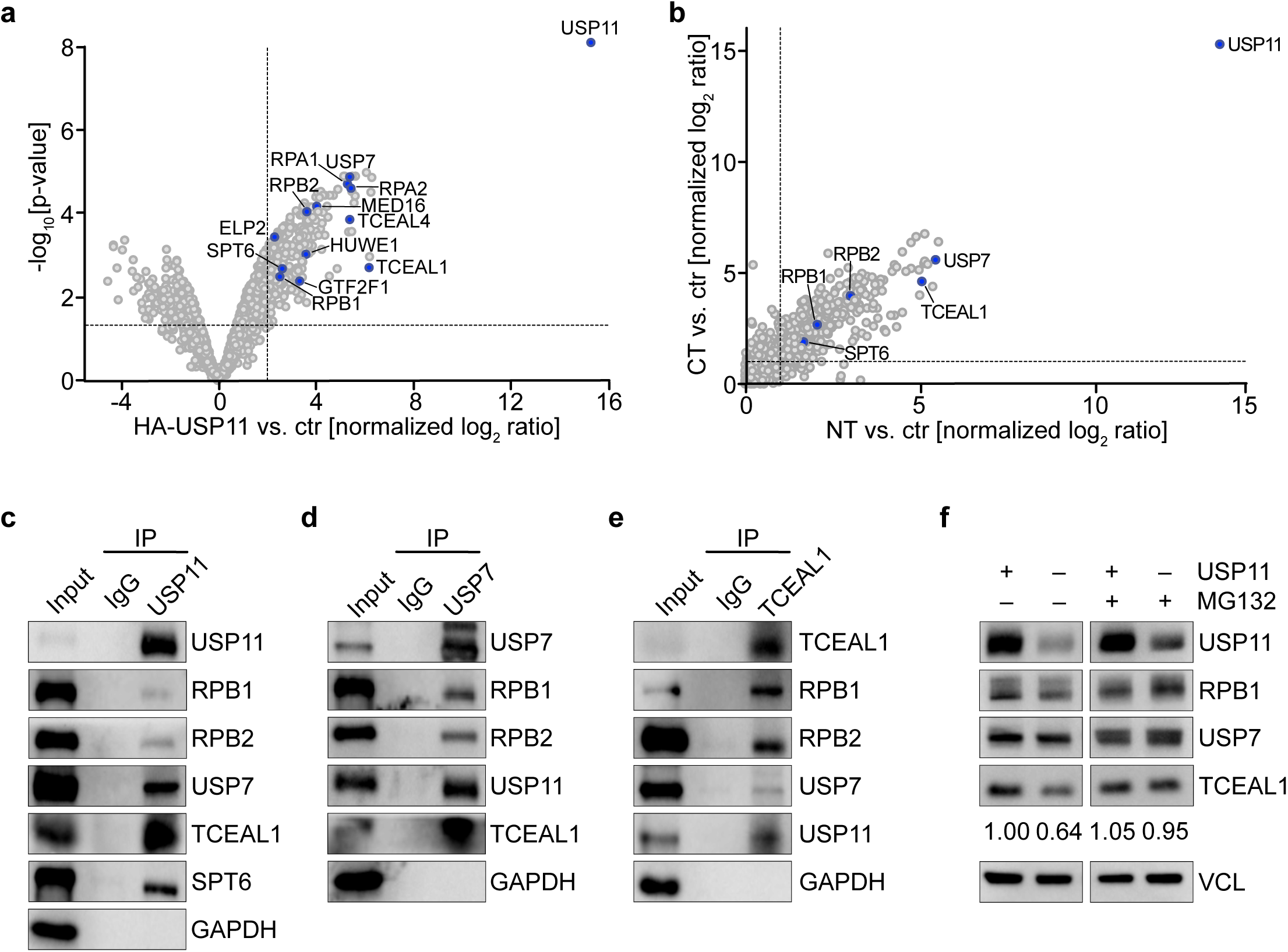
USP11 interacts with multiple proteins involved in basal transcription. **a.** Volcano plot of the consensus USP11 interactome with selected proteins labelled in blue. The x-axis displays the log_2_ ratio of proteins in HA-USP11 immunoprecipitations (IPs) relative to control IPs. The y-axis shows the -log_10_ of p-value for each protein. Vertical dashed line indicates log_2_ ratio of 2, horizontal dashed line indicates p-value of 0.05 (n=4; unless specified otherwise n indicates the number of independent biological replicates). **b.** Scatter plot of mass spectrometry data analysis of anti-HA IPs from SH-EP neuroblastoma cells stably expressing N-terminally (NT) or C-terminally (CT) HA-tagged USP11. The graph shows the normalized log_2_ ratio over anti-HA IP performed in vector control cells (ctr). Dashed lines indicate log_2_ ratio of 2 (n=2 for each IP). Proteins labeled in blue were validated in IPs shown below. **c.** Immunoblots of anti-USP11 IPs from SH-EP cells. The input corresponds to 1.5% of the amount used for the precipitation. Non-specific IgG and GAPDH were used as controls (n=2). **d.** Immunoblots of anti-USP7 IPs from SH-EP cells. The input corresponds to 1.5 % of the amount used for the precipitation. Non-specific IgG and GAPDH show controls (n=2). **e.** Immunoblots of anti-TCEAL1 IPs from SH-EP cells. The input corresponds to 2 % of the amount used for the precipitation. Non-specific IgG and GAPDH show controls (n=2). **f.** Immunoblots of indicated proteins in SH-EP cells upon knock-down of USP11 using inducible shRNA by adding doxycycline (1 µg/ml, 48 h). Cells were treated with MG132 (10 µM, 4 h) where indicated. VCL was used as a loading control (n=2).

To confirm these results, we immune-precipitated cell lysates using either USP11 or control antibodies and probed the precipitates with a series of antibodies against proteins that had been identified in the mass spectrometry. We confirmed the interactions of USP11 with the RNAPII subunits RPB1 and RBP2, the elongation factor SPT6, USP7 as well as with TCEAL1 (Figure 1c). Conversely, immunoprecipitation using an antibody against TCEAL1 immunoprecipitated RPB1 and RPB2, as well as USP7 and USP11 (Figure 1e). Finally, we showed that USP7 co-immunoprecipitates with RPB1, RPB2, USP11 and TCEAL1 (Figure 1d). Since USP11 is a deubiquitylating enzyme, we next tested whether USP11 stabilizes its associated proteins. We therefore depleted USP11 using shRNA and observed that USP11 silencing had little effect on USP7 and RPB1 levels but caused a moderate decrease in TCEAL1 levels (Figure 1f). The decrease in TCEAL1 levels upon USP11 depletion was abolished when cells were treated with the proteasome inhibitor MG132 arguing that TCEAL1 is stabilized by USP11 (Figure 1f). Collectively, the data show that USP11 associates with USP7, multiple proteins involved in basal transcription as well as TCEAL1 and that USP11 stabilizes TCEAL1.

### USP11, USP7 and TCEAL1 form a ternary complex in cells and in vitro

To determine whether USP11 forms a complex with USP7 and TCEAL1, we depleted TCEAL1 via stable expression of shRNAs in neuroblastoma cells and performed anti-USP11 immunoprecipitations (Supplemental Figure S2a). For these experiments, we used SH-EP MYCN-ER cells, that express a conditional MYCN-ER chimeric protein. In these cells, addition of hydroxy-tamoxifen (4-OHT) activates MYCN and enhances overall transcription elongation (Herold et al. 2019). These experiments showed that depletion of TCEAL1 strongly reduced binding USP11 to USP7 (Figure 2a). We also observed that USP11 associates with RNAPII and that depletion of TCEAL1 moderately reduced the association of both proteins. Next, we specifically probed for the association with pS2-RNAPII, which reflects the elongating form of RNAPII. Activation of MYCN increased levels of TCEAL1 and the association of USP11 with TCEAL1 and, to a lesser degree, with USP7. Importantly, USP11 was associated with pS2-RNAPII and depletion of TCEAL1 diminished the association of USP11 with pS2-RNAPII, arguing that TCEAL1 can stabilize complexes of USP11 with both total and elongating RNAPII (Figure 2a).

**Figure 2:**
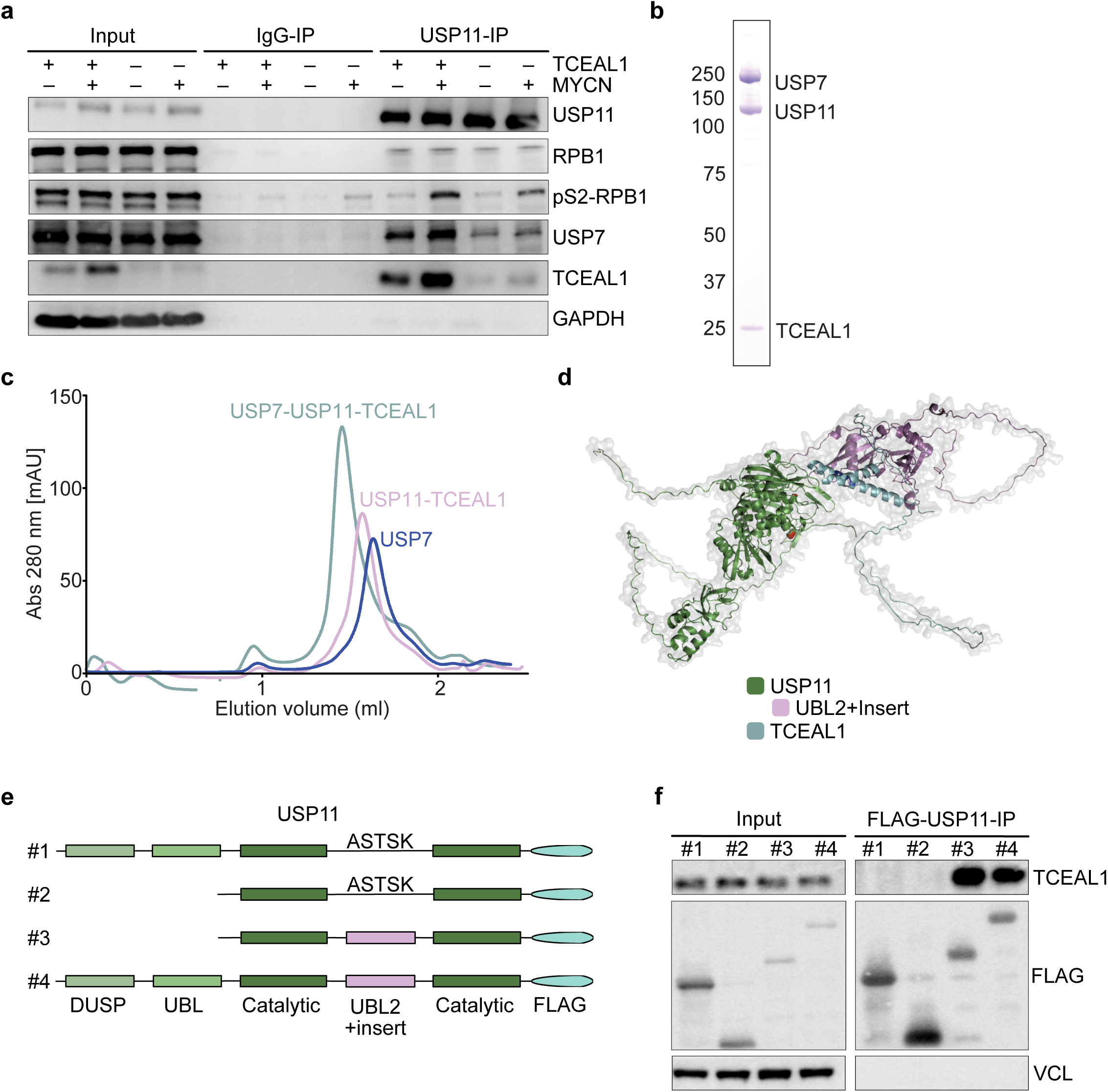
USP11 forms a ternary complex with USP7 and TCEAL1. **a.** Immunoblots of anti-USP11 IPs from SH-EP-MYCN-ER cells expressing doxycycline-inducible shRNA targeting *TCEAL1* in the presence or absence of doxycycline (1 µg/ml; 48 h) and MYCN activation (200 nM 4-OHT, 4 h). The input corresponds to 1.5 % of the amount used for the precipitation. Non-specific IgG and GAPDH were used as controls (n=3). **b.** Coomassie blue-stained 4–12 % SDS–PAGE gel of purified USP11, TCEAL1 and USP7. **c.** The indicated recombinant proteins were incubated with each other and resulting complexes were separated by size-exclusion chromatography. Curves show milli absorption (Abs) units (mAU) at 280 nm (n=2). **d.** AlphaFold2 prediction of USP11/TCEAL1 complex structure (highest confidence model of five is shown). **e.** Schematic diagram of FLAG-tagged USP11 deletion mutant proteins. **f.** Immunoblots of indicated proteins from pulldown assays of SH-EP cell lysates incubated with recombinant FLAG-tagged USP11 alleles. VCL was used as loading control (n=2).

To test whether USP11, USP7 and TCEAL1 directly form a ternary complex, we recombinantly expressed and purified the three proteins and performed size exclusion chromatography on a USP11-TCEAL1 complex, isolated USP7, and a mixture containing the USP11-TCEAL1 complex and USP7. This showed that USP11-TCEAL1 and USP7 elute in later fractions than when all three proteins were co-incubated, indicating that USP11, TCEAL1 and USP7 indeed form a stable ternary complex (Figure 2b,c). To understand the assembly of the binary USP11-TCEAL1 complex, we used AlphaFold2 (Jumper et al. 2021; Varadi et al. 2022) to model the interaction between USP11 and TCEAL1. Several models of the complex predicted that a domain inserted into the catalytic domain of USP11 is important for the interaction (Figure 2d and Supplemental Figure S2b). This domain comprises of an ubiquitin-like (UBL) domain with an insert and is hence termed the “UBL2+insert” domain of USP11 (Harper et al. 2014). To test this prediction, we expressed a small series of deletion mutants of USP11 as FLAG-tagged proteins in cells and performed *in vivo* pulldown assays (Figure 2e). These confirmed that the UBL2+insert domain of USP11 is critical for the interaction with TCEAL1 (Figure 2f). Collectively, the data argue that USP11, USP7 and TCEAL1 form a ternary complex and that this complex is associated with RNAPII.

### TCEAL1 globally binds to active promoters via its TFIIS-homology domain

To investigate whether TCEAL1 associates with chromatin, we stably expressed HA-tagged TCEAL1 and performed chromatin-immunoprecipitation and sequencing (ChIP-seq) experiments. As negative control we performed HA-immunoprecipitations from cells that did not express HA-TCEAL1. Visual inspection of browser tracks of individual genes showed that TCEAL1 associated with chromatin surrounding the transcription start sites (TSS) of all inspected actively transcribed genes and that its binding sites largely overlapped with those of total RNAPII close to the TSS (Figure 3a). Genome-wide analyses confirmed this observation and showed that the amount of TCEAL1 bound to each promoter paralleled the amount of promoter-bound RNAPII across all active genes (Figure 3b). The homology between TCEAL1 and TFIIS is restricted to a specific sequence (<u>T</u>FIIS-<u>h</u>omology <u>s</u>equence, THS) that in TFIIS is part of domain II and forms an alpha-helix which is involved in the interaction surface with RNAPII (Figure 3c) (Hijazi et al. 2022). To test whether the THS in TCEAL1 is required for chromatin association and binding to RNAPII, we stably expressed different HA-tagged TCEAL1 mutants: wild-type (TCEAL1^wt^), one lacking the alpha-helix (TCEAL^λ-.116-139^ ^aa^) or alleles with mutations that alter either three (D130R, E131R, L132A; generating TCEAL^3RA^) or five (D130R, E131R, L132A, E133R, E134R; generating TCEAL^5RA^) amino acids and cause a major change in its charge (Figure 3d). Immunoblots showed that deletion of the THS compromised TCEAL1 stability, whereas the point-mutated alleles were expressed at the same level as the wildtype protein (Figure 3e). ChIP experiments followed by qPCR showed that all three mutant variants were strongly compromised in their ability to associate with promoters (Figure 3f). Collectively the data show that TCEAL1 globally associates with active promoters and that the THS is required for chromatin association.

**Figure 3:**
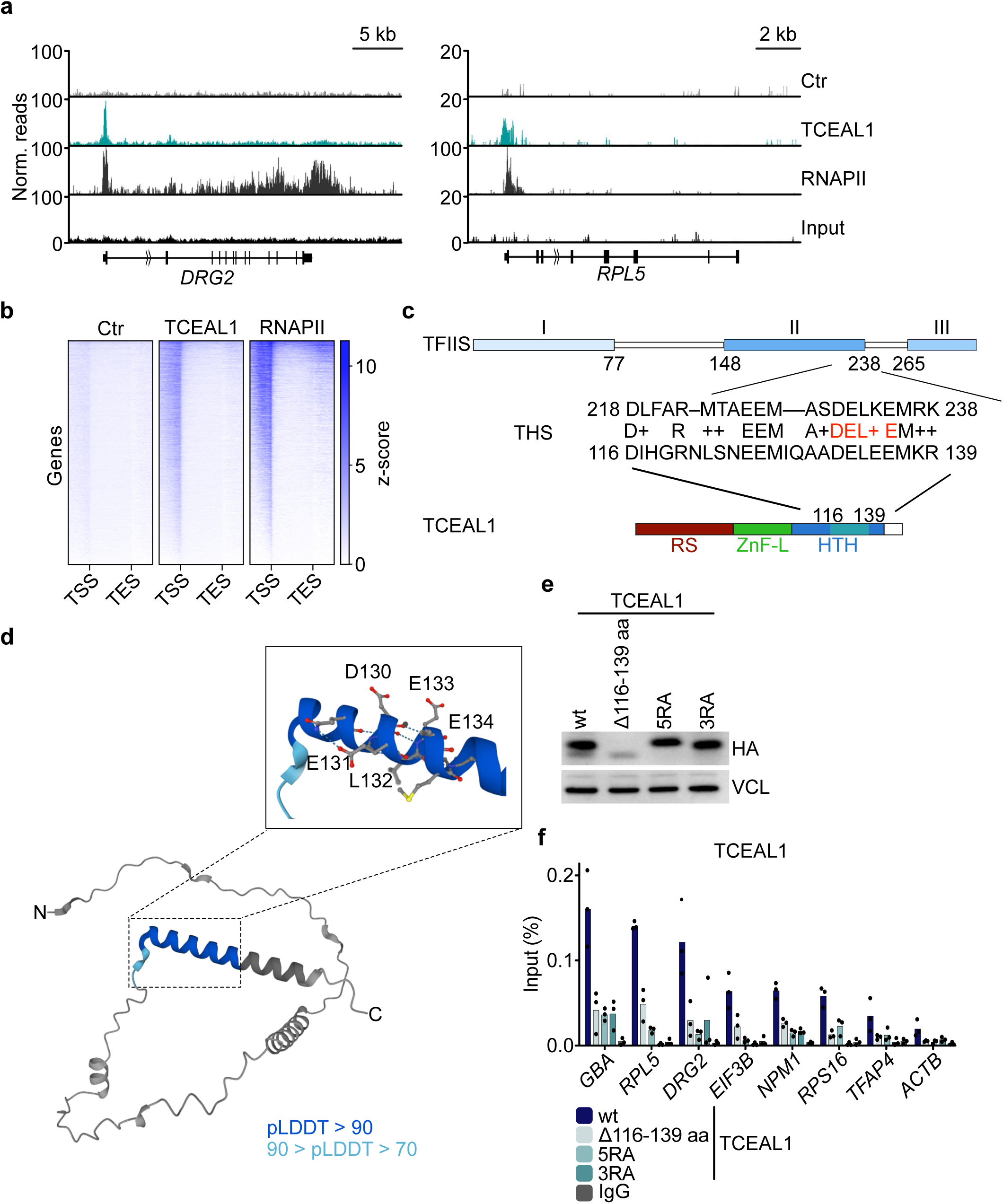
Chromatin association of TCEAL1 depends on the TFIIS homology region. **a.** Genome browser tracks of indicated loci showing chromatin association of HA-TCEAL1 and RNAPII in SH-EP cells. anti-HA chromatin immunoprecipitations (ChIPs) from cells that do not express HA-TCEAL1 were used as control (ctr) (n=2). **b.** Heatmaps indicating global chromatin occupancy of TCEAL1 and RNAPII at active genes. The heatmap is sorted according to RNAPII binding (TSS, transcription start site; TES, transcription end site, N = 16,285 genes analyzed). **c.** Sequence alignment of TFIIS and TCEAL1, with the domains of the TFIIS protein shown on top and of TCEAL1 shown below. The turquoise color in the HTH indicates the homology domain with TFIIS (THS) (RS, Arginine/Serine rich domain; ZnF-L, Zinc-Finger like domain; HTH, Helix-Turn-Helix domain). Amino acids subjected to mutagenesis are highlighted in red. **d.** AlphaFold2 prediction of TCEAL1 structure. Predicted local distance difference test (pLDDT) indicates modeling confidence and is marked by different shades of blue. Zoom-in: alpha-helix conserved between TCEAL1 and TFIIS. Amino acids subjected to mutagenesis are labelled. **e.** Immunoblots of SH-EP cells expressing HA-tagged TCEAL1 wildtype or the indicated mutant alleles. VCL was used as a loading control (n=2). **f.** Anti-HA ChIP at indicated promoter regions in SH-EP cells expressing HA-tagged TCEAL1 wildtype or the indicated mutant alleles. IgG was used as negative control. Shown is the mean of technical replicates of one representative experiment (n=3).

### TCEAL1 antagonizes TFIIS binding to RNAPII and chromatin

To explore the hypothesis that TCEAL1 antagonizes the access of TFIIS to RNAPII, we generated cells with stably expressing doxycycline-inducible shRNA targeting *TCEAL1* (Supplemental Figure S3a) and performed TFIIS ChIP-qPCR experiments and spike normalized TFIIS ChIP-seq (ChIP-Rx). Upon depletion of TCEAL1, more TFIIS was bound at the TSS of multiple individual promoters (Figure 4a,b). Genome-wide analyses confirmed that TFIIS chromatin association increased globally at active promoters upon depletion of TCEAL1 (Figure 4c,d and Supplemental Figure S3b). To discern the status of promoter-proximal RNAPII on which TCEAL1 acts, we performed proximity ligation assays (PLAs, Figure 4e,f and Supplemental Figure S3c). PLAs showed that depletion of TCEAL1 caused a mild increase in the proximity of TFIIS to total RNAPII and a stronger and statistically significant with RNAPII that is phosphorylated at Ser5 in the CTD (pS5-RNAPII), which mainly reflects promoter-proximal active RNAPII (Hsin and Manley 2012). In contrast, depletion of TCEAL1 caused a mild decrease in the PLA signal between TFIIS and non-phosphorylated RNAPII (Figure 4e,f). To explore whether TCEAL1 exerts these effects by competing with TFIIS for access to RNAPII, we stably expressed either TCEAL1^wt^ or the TCEAL1^3RA^ allele that shows reduced association with chromatin and measured both, chromatin association of TFIIS and binding of TFIIS to RNAPII. Neither the expression of TCEAL1^wt^ nor of TCEAL1^3RA^ had an effect on TFIIS protein levels (Figure 4g). ChIP-Rx showed that ectopic expression of TCEAL1^wt^ globally decreased TFIIS association with active promoters (Figure 4h, and Supplemental Figure S3d). In contrast, TCEAL1^3RA^ enhanced global chromatin association of TFIIS (Figure 4h and Supplemental Figure S3d). Consistently, immunoprecipitation assays showed that expression of TCEAL1^wt^ reduced the association of TFIIS with RNAPII while expression of TCEAL1^3RA^ did not (Figure 4g). We concluded that TCEAL1 antagonizes chromatin association and binding of TFIIS to RNAPII in a THS-dependent manner, arguingthat these effects are due to direct competition between both proteins.

**Figure 4:**
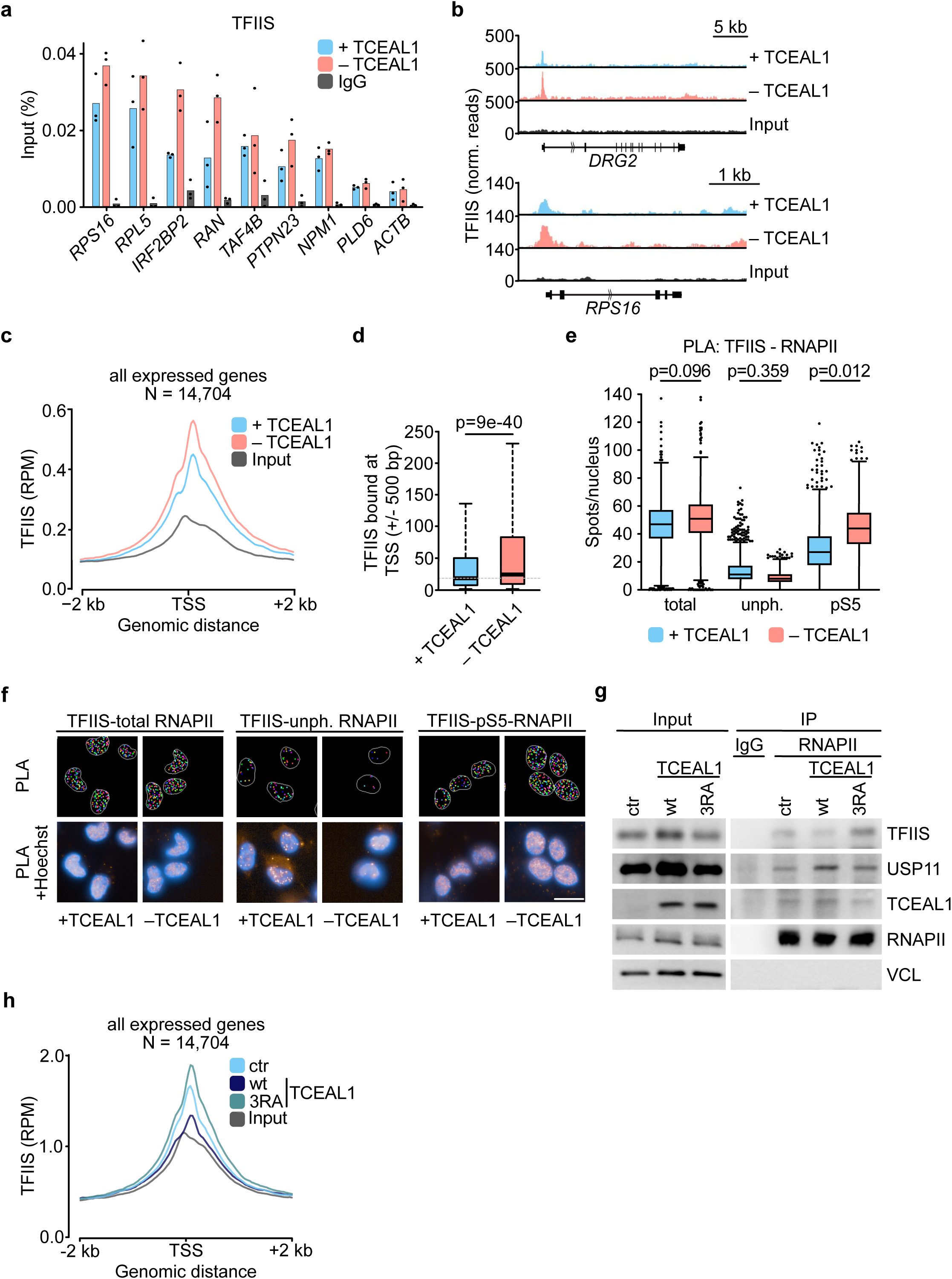
TCEAL1 and TFIIS compete for binding to RNAPII. **a.** TFIIS ChIP at indicated loci in SH-EP cells expressing a doxycycline-inducible shRNA targeting *TCEAL1* (1 µg/ml; 48 h). IgG was used as negative control. Shown is the mean of technical triplicates of one representative experiment (n=3). **b.** Genome browser tracks of indicated loci showing TFIIS spike-in ChIP (ChIP-Rx) data in cells as described in a (n=2). **c.** Global average read density of TFIIS ChIP-Rx at the TSS (N = genes analyzed). **d.** Boxplot representing global reads of TFIIS at the TSS upon TCEAL1 knock-down. Grey line indicates mean of unperturbed cells. P-value was calculated using a paired two-sided Student’s t-test with unequal variance (N = 21,005 genes were analyzed). **e.** Boxplot of single-cell analysis of nuclear proximity ligation assay (PLA) foci between TFIIS and differently phosphorylated RNAPII of one representative PLA experiment. P-values were calculated using an unpaired t-test (n=3). **f.** Representative PLA images showing detected PLA spots (top) and PLA signal merged with Hoechst33342 staining (below). Scale bar indicates 20 µm (n=3). **g.** Immunoblots of RNAPII IPs of SH-EP cells expressing HA-tagged TCEAL1^wt^ or TCEAL1^3RA^. Co-precipitated proteins are indicated. Non-specific IgG was used as control (n=1). **h.** Global average read density of TFIIS ChIP-seq data in cells upon stable expression of TCEAL1^wt^ or TCEAL1^3RA^ (N = genes analyzed, n=2).

### USP11 stabilizes RPB8, a core subunit of RNAPII

To explore a potential role of USP11 during early transcription, we performed mass spectrometry of cells measuring protein abundance before and after depletion of USP11. This revealed that USP11 depletion caused a decrease in steady-state levels of a small group of proteins that includes RPB8, which was confirmed by immunoblots (Figure 5a,b). RPB8 is a common subunit of all three nuclear RNA polymerases and is a target for BRCA1-mediated ubiquitylation upon DNA damage (Wu et al. 2007). The mass spectrometry data also showed that depletion of USP11 caused weaker decreases in the levels of multiple other subunits of nuclear RNA polymerases (Figure 5c,d). Using quantitative immunofluorescence, we confirmed that USP11 depletion reduced RPB8 abundance in the nucleoplasm and potentially also in the nucleolus, while it had no effect on the levels of RPB1, another subunit of RNAPII (Figure 5e).

**Figure 5:**
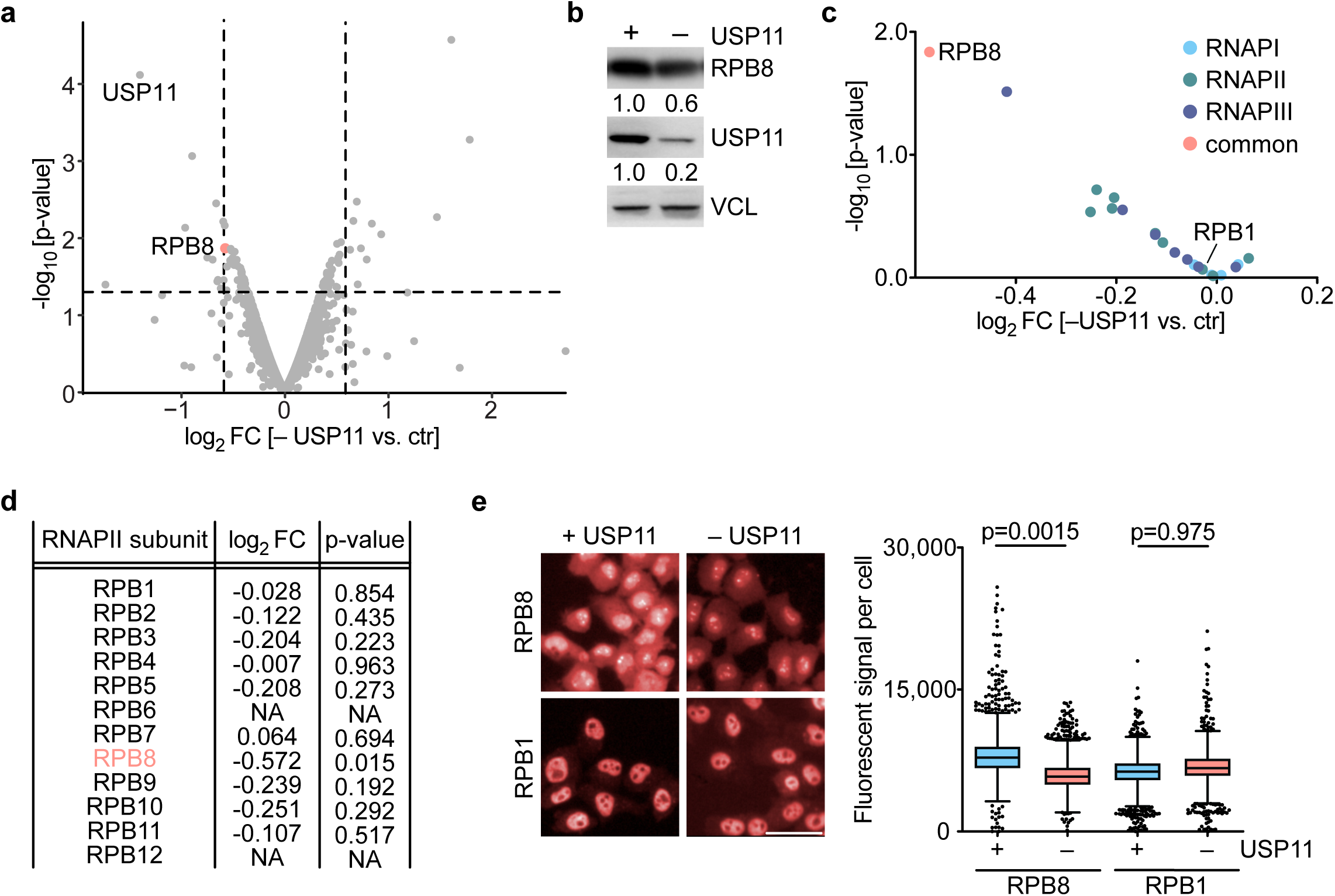
USP11 stabilizes RPB8. **a.** Volcano plot of mass spectrometry data of IMR-5 cells expressing a doxycycline-inducible shRNA targeting *USP11* (48 h, 1 µg/ml). The x-axis displays log_2_FC of proteins in cells with control levels (ctr) or knock-down of USP11 (– USP11). The y-axis shows the -log_10_ p-value for each protein (n=2). **b.** Immunoblots from cells described in a. VCL was used as loading control (n=3). **c.** Volcano plot showing changes of different RNAP subunits after knock-down of USP11. Axis as described in a. **d.** Table showing log_2_FC of different RNAPII subunits after shRNA-mediated depletion of USP11. **e.** Representative images of SH-EP cells immuno-stained for RPB8 or RPB1 after knock-down of USP11. Scale bar indicates 50 µM (left). Boxplots of single-cell analysis of a representative experiment (right). P-values were calculated using an unpaired t-test (n=3).

### TCEAL1 stabilizes RNAPII on chromatin to sustains a mesenchymal expression program

To understand whether TCEAL1 affects RNAPII function in cells, we initially performed chromatin-immunoprecipitations under different conditions. We noted a strong increase in the association of TCEAL1 with several promoters when cells had been treated with flavopiridol, a potent inhibitor of CDK9 that promotes pause release of RNAPII (Supplemental Figure S4a). Since flavopiridol also inhibits other kinases, we performed ChIP-seq experiments from cells treated with either THZ1, a selective inhibitor of the CDK7 kinase (Kwiatkowski et al. 2014), or with NVP-2, a specific inhibitor of the CDK9 kinase (Olson et al. 2018). These experiments showed that inhibition of CDK7 had little effect on the association of TCEAL1 with active promoters, while inhibition of CDK9 increased the chromatin association of TCEAL1 and shifted the peak of association further downstream to a position corresponding to that of paused RNAPII (Figure 6a).

**Figure 6:**
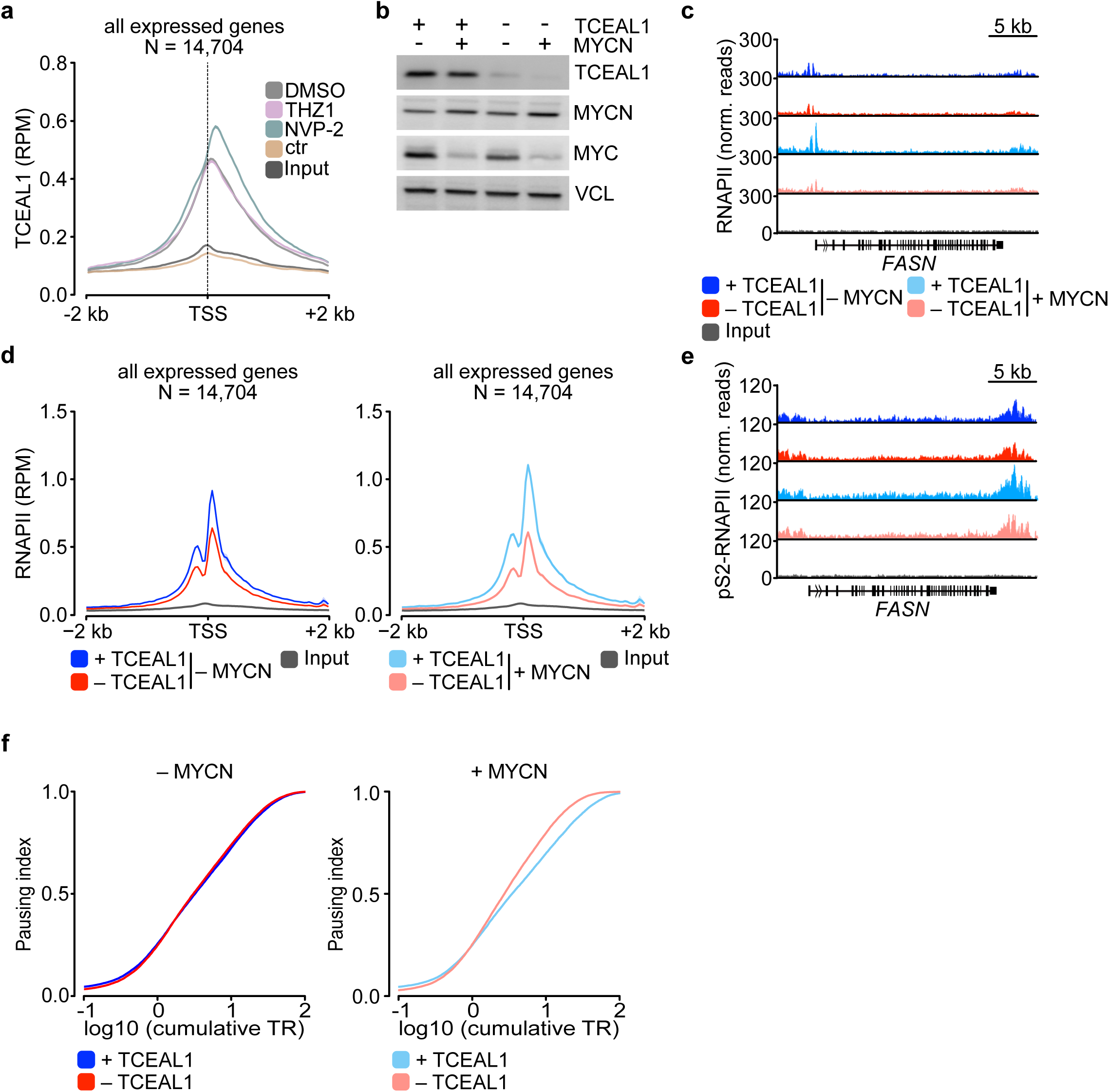
TCEAL1 stabilizes RNAPII during early transcription. **a.** Global average read density plot of HA ChIP-seq data in SH-EP cells expressing HA-TCEAL1 after treatment with THZ1 (200 nM, 8 h), NVP-2 (1 µM, 3 h) or DMSO as control. SH-EP cells were used as negative control (ctr). Data show mean ± SEM (N = genes analyzed, n=2). **b.** Immunoblots of indicated proteins in SH-EP-MYCN-ER cells expressing doxycycline-inducible shRNA targeting *TCEAL1* (Dox: 1 µg/ml; 48 h) and MYCN activation (200 nM, 4 h 4-OHT). VCL was used as a loading control (n=3). **c.** Genome browser tracks of the *FASN* locus showing chromatin association of RNAPII in SH-EP-MYCN-ER cells (treatment as in b) in the presence (blue) or after knock-down (red) of TCEAL1 without (top) or with (bottom) MYCN induction. **d.** Global average read density of RNAPII ChIP-Rx data as described in c. Data show mean ± SEM (N = genes analyzed, n=3). **e.** Genome browser tracks of the *FASN* locus showing chromatin association of pS2-RNAPII in in SH-EP-MYCN-ER cells as described in b (n=2). **f.** Pausing index of RNAPII. Ratios were calculated for cells described in panel b, either without (left) or with (right) MYCN induction (N = 22,725 genes analyzed).

To determine whether TCEAL1 affects chromatin association of RNAPII, we depleted TCEAL1 using specific shRNAs and performed ChIP-Rx using antibodies that recognize total RNAPII and elongating pS2-RNAPII. To better discern the effects on overall chromatin binding and elongation, we performed these experiments in SH-EP-MYCN-ER cells (Figure 6b). Activation of MYCN suppresses expression of endogenous MYC in these cells (Herold et al. 2019). Notably, depletion of TCEAL1 did not affect levels of either protein, although USP11 and USP7 can stabilize MYC and MYCN, respectively (Tavana et al. 2016; Pornour et al. 2024). This argues that effects of TCEAL1 are not mediated indirectly by effects on MYC and MYCN levels and also suggests that formation of ternary complex may not be required for all functions of USP11.

Inspection of multiple individual promoters and genome-wide analyses showed that chromatin association of RNAPII was strongly reduced upon depletion of TCEAL1 (Figure 6c,d and Supplemental Figure S4b). ChIP-Rx using an antibody which recognizes pS2-RNAPII showed a decrease in chromatin association that paralleled the decrease in total RNAPII (Figure 6e). Analysis of the pausing index, that indicates the fraction of RNAPII bound to the promoter relative to that present in gene bodies, showed no difference upon depletion of TCEAL1, arguing that the decrease in promoter association of RNAPII is not a consequence of decreased pausing or enhanced elongation (Figure 6f). Consistently, depletion of TCEAL1 also strongly reduced chromatin association of RNAPII in cells in which the MYCN-ER chimera had been acutely activated (Figure 6d).

To understand whether the stabilization of RNAPII by TCEAL1 is particularly relevant for a specific group of genes, we performed RNA sequencing upon TCEAL1 depletion. Gene set enrichment analyses (GSEA) showed that genes that are induced during the TGF-beta dependent transition from epithelial to mesenchymal cells were highly enriched among TCEAL1-dependent genes (Figure 7a,b and Supplemental Figure S5a). In multiple tumor entities, this gene expression program is associated with invasive and metastatic tumors and consistently, gene sets that define such tumors were significantly downregulated upon depletion of TCEAL1 (Figure 7b). Neuroblastoma tumors harbor two major histological cell types, adrenergic and mesenchymal cells, which differ in their gene expression pattern, biology, clinical features and response to therapy (van Groningen et al. 2017). Expression of most TCEAL1-dependent genes was positively correlated with a mesenchymal and anticorrelated with an adrenergic gene expression program in large gene expression databases of primary neuroblastoma patients (Figure 7c). Overall, the TCEAL1-dependent gene was clearly correlated with mesenchymal cells both in a panel of neuroblastoma cell lines (Figure 7d) as well as in primary tumors. Analysis of ChIP-seq data showed that both TCEAL1 and TFIIS were bound to the promoters of key genes that define the mesenchymal gene expression including THBS1, encoding thrombospondin, and TGFB1, encoding TGF beta 1 (Supplemental Figure S5b). Depletion of TCEAL1 strongly reduced RNAPII occupancy and enhanced TFIIS occupancy at these promoters and multiple other mesenchymal genes (Supplemental Figure S5b). We concluded that TCEAL1 directly sustains a mesenchymal gene expression program in neuroblastoma and potentially other solid tumors.

**Figure 7:**
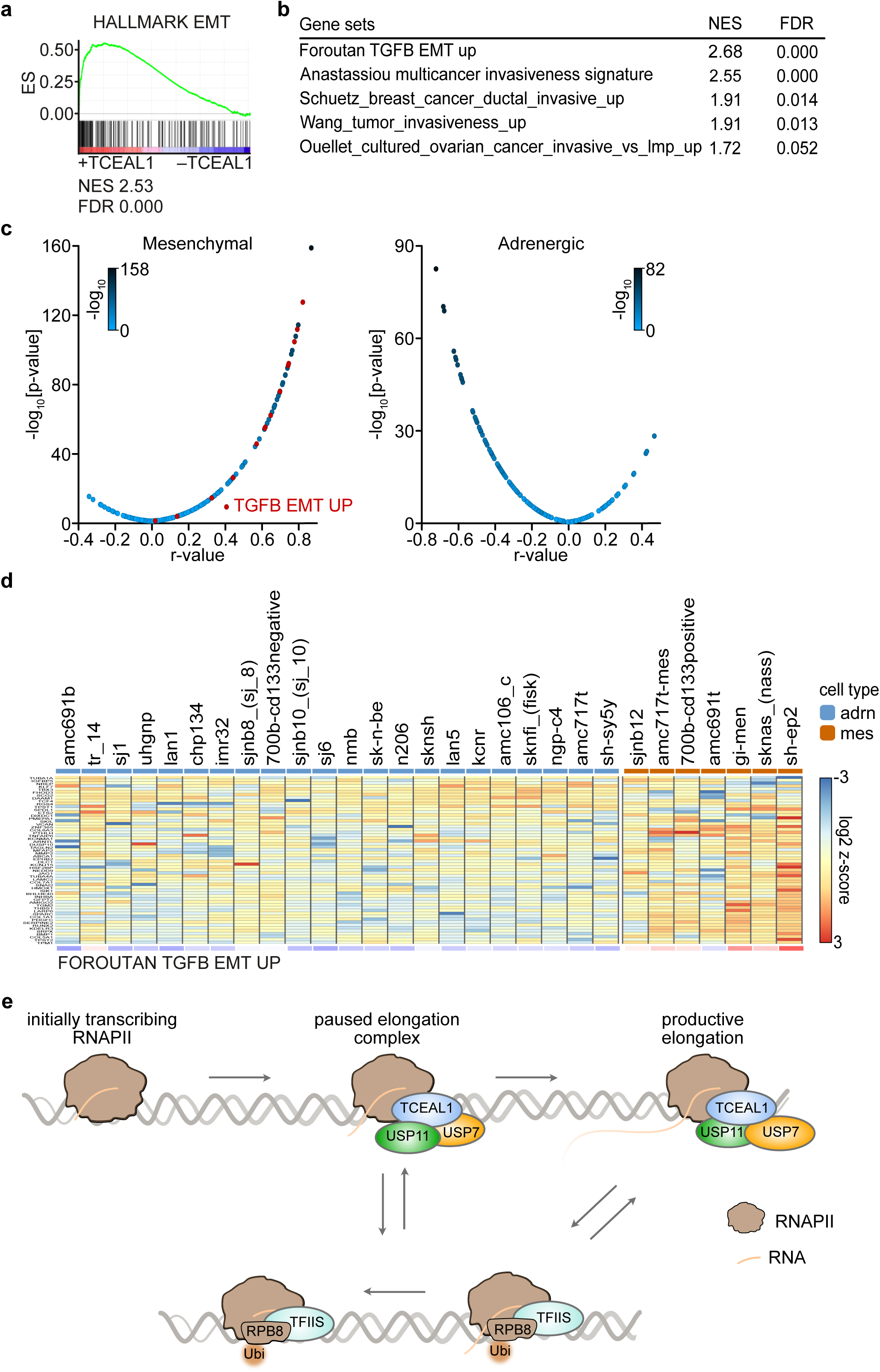
TCEAL1 promotes a mesenchymal gene expression program. **a.** GSEA plot of the hallmark gene set “epithelial mesenchymal transition” upon TCEAL1 depletion in SH-EP cells. NES, normalized enrichment score; FDR, false discovery rate. **b.** Table of additional gene sets associated with the TGF beta-dependent epithelial-mesenchymal transition of breast cancer cells and metastasis that are downregulated upon depletion of TCEAL1. **c.** Correlation of the expression of the most strongly TCEAL1-dependent target genes with mesenchymal (left) and adrenergic (right) gene expression program in n=498 primary neuroblastoma samples. N = 100 genes were analyzed in each case. Genes contained in an TGF-beta dependent EMT gene set (Foroutan, see panel b) are indicated in red. **d.** Heatmap comparing adrenergic (blue) and mesenchymal (orange) cells to the most significant downregulated genes upon depletion of TCEAL1 depletion. R-values comparing expression of genes of each cell line cell lines to Foroutan signature are depicted at the bottom. **e.** Proposed Model illustrating that the USP11/TCEAL1/USP7 complex stabilizes RNAPII during early transcription both by buffering TFIIS activity and via deubiquitylation of RBP8.

## Discussion

Here we investigated the function of USP11, a deubiquitylating enzyme that has been implicated in transcription-related processes and in the maintenance of genome stability (Orthwein et al. 2015; Deng et al. 2018; Herold et al. 2019; Jurga et al. 2021; Pornour et al. 2024), by analyzing its interactome in human neuroblastoma cells. We found that USP11 together with its heterodimeric partner USP7, forms a trimeric complex with TCEAL1, a member of the TCEAL family of proteins, and that TCEAL1 stabilizes the USP11/USP7 heterodimer. TCEAL1 interacts with RNAPII via a short helix that shares sequence homology with a region in TFIIS. TCEAL1 and USP11 stabilize RNAPII during early transcription by competing with TFIIS and by protecting RPB8, an essential subunit of RNAPII, from degradation, suggesting that TCEAL1 and USP11 maintain RNAPII in a state that is poised for elongation (Figure 7e). RNA sequencing shows that a specific group of downstream target genes most clearly depends on this mechanism, which comprises of a TGF beta-dependent gene expression program.

Multiple previous observations have linked USP11 to the TGF beta pathway since it deubiquitylates and stabilizes the TGF beta receptor (Al-Salihi et al. 2012; Jacko et al. 2016). As consequence, USP11 maintains a range of cell types in an TGF beta-responsive state (Istomine et al. 2019). Maintaining RNAPII in a poised state, as reported here, is likely to contribute to this responsiveness. The TCEAL1-dependent gene expression program is very similar to TGF beta-dependent gene expression programs observed in breast, colon and hepatocellular carcinoma during the epithelial-mesenchymal transition and, as a result, correlates with invasiveness and metastasis. Indeed, USP11 expression is upregulated during and is essential for EMT. Depletion of USP11 reduces the clonogenic growth of breast cancer cells and the number of pulmonary metastases (Garcia et al. 2018). Similarly, USP11 has been causally implicated in the metastasis of colon and hepatocellular carcinoma via its effect on TGF beta signaling and additional downstream targets (Zhang et al. 2018; Sun et al. 2019; Huang et al. 2021; Qiao et al. 2024). Here we show that the TCEAL1-dependent gene expression program is highly correlated with mesenchymal, but not adrenergic neuroblastomas (van Groningen et al. 2017). Importantly, mesenchymal tumor cells show enhanced resistance to immunotherapy and targeted therapy using ALK kinase inhibitors (Mabe et al. 2022; Sengupta et al. 2022; Westerhout et al. 2022) hence inhibiting the underlying gene expression program may improve responses to both therapies.

Considering the widespread reduction of RNAPII at promoters, it is surprising that a specific group of genes responds most strongly to TCEAL1 depletion. The reasons for this specificity are unknown at this point, but TGF beta-dependent gene expression is known to depend on KAP1/TRIM28, a multidomain protein, which binds acetylated histones, recruits BRD4 to promoters and facilitates SMAD-dependent CDK9 recruitment and transcription elongation (Bacon et al. 2020). TRIM28/KAP1 is a member of a family of four related RING finger ubiquitin ligases that can trigger the ubiquitylation and degradation of target proteins; indeed, three of them, TRIM24, TRIM28 and TRIM33 associate with each other (Herquel et al. 2011). Intriguingly, TRIM33 ubiquitylates SMAD proteins and the TGF beta receptor and the complex of the three TRIM proteins suppresses the development of hepatocellular carcinomas (Dupont et al. 2009; Herquel et al. 2011; Quere et al. 2014). USP11 interacts with TRIM28/KAP1 (Perry et al. 2021; Jin et al. 2022) and we hypothesize that this interaction negates the ubiquitin ligase activity of TRIM28/KAP1 while maintaining its ability to bind acetylated histones and recruit BRD4, and that KAP1/TRIM28 and TCEAL1/USP7/USP11 therefore cooperate to promote TGF beta-dependent gene expression.

Collectively, our data show TCEAL1/USP11 complexes alter RNAPII at core promoters to be responsive to specific inducing signals that are known to control tumor progression and therapy resistance. Since the physiological function of TCEAL1 is restricted to CNS development and TCEAL1 may be largely dispensable in adult organisms, the findings also suggest that interference with the function of the TCEAL1/USP11/USP7 complex using small molecules is a valid therapeutic strategy to abrogate a specific gene expression program that drives therapy resistance, tumor cell invasion and metastasis.

## Materials and Methods

### Cell culture

Neuroblastoma cell lines (SH-EP and IMR-5) were grown in RPMI 1640 medium (Thermo Fisher Scientific). HEK293TN and NIH-3T3 cells were grown in DMEM (Thermo Fisher Scientific). Medium was supplemented with 10 % fetal calf serum (Biochrom and Sigma-Aldrich) and penicillin-streptomycin (Sigma-Aldrich). All cells were routinely tested for mycoplasma contamination. Where indicated, cells were treated with 4-OHT (Sigma-Aldrich), MG132 (Calbiochem/Merck), NVP-2 (Tocris Biosciences), Flavopiridol (Sigma-Aldrich), THZ1 (Hycultec) or DMSO (Sigma-Aldrich).

### Cloning

Constructs to generate shRNAs targeting the proteins USP11 or TCEAL1 were cloned into a lentiviral pLT3GEPIR or pGIPZ vector, which harbor a miR-E based backbone based on the design reported initially (Fellmann et al. 2013). For overexpression studies double-stranded DNA fragments (gBlock™ User Guide, IDT) based on cDNA sequences of USP11 and TCEAL1 were cloned into a pLEGO or pRRL vector containing an SFFV promoter. To engineer proteins with loss or gain of function, site directed mutagenesis (SDM) was performed. Mutations were integrated by PCR using pairs of oligonucleotide primers designed mismatching nucleotides with the coding codon of interest. For transformation of plasmids, XL-1 blue *E. coli* cells were used. The integrity of the DNA was confirmed by Sanger sequencing (LGC genomics) of the expression plasmids containing the inserts of interest.

### Transfection and lentiviral infection

Transfection with cDNA was performed using PEI (Sigma-Aldrich). Cells were collected 48 h after transfection. For lentivirus production, HEK293TN cells were transfected using PEI. Lentivirus expressing plasmids were transfected together with the packaging plasmid psPAX.2 and the envelope plasmid pMD2.G. Virus-containing supernatant was collected 48 h and 72 h after transfection. SH-EP and IMR-5 cells were infected with lentiviral supernatant in the presence of 4 µg/ml polybrene for 24 h. Cells were selected for 2 days with puromycin (SH-EP: 2 µg/ml, IMR-5: 0.5 µg/ml) and afterwards seeded for the experiment.

### Immunoblot

Cells were lysed in RIPA buffer (50 mM HEPES pH 7.9, 140 mM NaCl, 1 mM EDTA, 1 % Triton X-100, 0.1 % SDS, 0.1 % sodium deoxycholate) containing protease and phosphatase inhibitors (Sigma-Aldrich) and incubated for 20 min at 4 °C. The lysate was cleared by centrifugation and the protein concentration was determined using a BCA assay. Cell lysate was separated by Bis-Tris-PAGE and transferred to PVDF membranes (Millipore). Membranes were blocked for 1 h and incubated using the indicated antibodies overnight at 4 °C. Antibodies used are listed in Table S1. As loading control, vinculin (VCL), actin (ACTB) or GAPDH was used. Afterwards, membranes were washed and probed for 1 h at RT with HRP-conjugated secondary antibodies. Images were acquired using LAS3000 or LAS4000 Mini Image System (Fuji).

### Immunoprecipitation

For co-immunoprecipitations, 20 µl per IP of 1:1 A/G Dynabeads mix (Thermo Fisher Scientific) was washed with 5 mg/ml BSA-PBS and then incubated overnight at 4 °C on a rotating wheel with 2.5 µg antibody against the protein of interest diluted in BSA-PBS. Next, cell pellets were lysed with HEPES lysis buffer (20 mM HEPES pH 7.9, 150 mM NaCl, 0.2 % v/v NP-40, 0.5 mM EDTA, 10 % v/v Glycerol, 2 mM MgCl_2_) and incubated with 50 U Benzonase (Merck) for 1 h at 4 °C on a rotating wheel. 2-4 mg of lysate were added to the bead/antibody mix and incubated for 6 h at 4 °C on a rotating wheel. 1.5-2 % of the lysate was kept as input reference. After thoroughly washing with HEPES lysis buffer the elution was carried out by resuspending the beads in 2x Laemmli buffer and boiling for 5 min at 95 °C. The samples were then subjected to immunoblotting.

### NanoLC-MS/MS analysis of USP11 interactors

Protein precipitation was performed overnight at -20 °C with a fourfold volume of acetone. Pellets were washed with acetone at -20 °C. Precipitated proteins were dissolved in NuPAGE® LDS sample buffer (Life Technologies), reduced with 50 mM DTT at 70 °C for 10 min and alkylated with 120 mM Iodoacetamide at RT for 20 min. Separation was performed on NuPAGE® Novex® 4-12 % Bis-Tris gels (Life Technologies) with MOPS buffer according to manufacturer’s instructions. Gels were washed and stained for 1 h with Simply Blue™ Safe Stain (Life Technologies). After washing for 1 h, each gel lane was cut into 15 slices. The excised gel bands were destained with 30 % acetonitrile in 0.1 M NH_4_HCO_3_ (pH 8), shrunk with 100 % acetonitrile, and dried in a vacuum concentrator (Concentrator 5301, Eppendorf, Germany). Digests were performed with 0.1 µg trypsin (Trypsin Gold, Mass Spectrometry Grade, Promega) per gel band overnight at 37 °C in 0.1 M NH_4_HCO_3_ (pH 8). After removing the supernatant, peptides were extracted from the gel slices with 5 % formic acid, and extracted peptides were pooled with the supernatant. NanoLC-MS/MS analyses were performed on an Orbitrap Fusion (Thermo Scientific) equipped with a PicoView Ion Source (New Objective) and coupled to an EASY-nLC 1000 (Thermo Scientific). Peptides were loaded on capillary columns (PicoFrit, 30 cm x 150 µm ID, New Objective) self-packed with ReproSil-Pur 120 C18-AQ, 1.9 µm (Dr. Maisch) and separated with a 30-minute linear gradient from 3 % to 30 % acetonitrile and 0.1 % formic acid and a flow rate of 500 nl/min. Both MS and MS/MS scans were acquired in the Orbitrap analyzer with a resolution of 60,000 for MS scans and 15,000 for MS/MS scans. HCD fragmentation with 35 % normalized collision energy was applied. A Top Speed data-dependent MS/MS method with a fixed cycle time of 3 s was used. Dynamic exclusion was applied with a repeat count of 1 and an exclusion duration of 30 s; singly charged precursors were excluded from selection. The minimum signal threshold for precursor selection was set to 50,000. Predictive AGC was used with AGC a target value of 2e5 for MS scans and 5e4 for MS/MS scans. EASY-IC was used for internal calibration. Raw MS data files were analyzed with MaxQuant version 1.6.2.2 (Cox and Mann 2008). The database search was performed with Andromeda, which is integrated in the utilized version of MaxQuant. The search was performed against the UniProt human database (June 27,2018, UP000005640, 73,099 entries). Additionally, a database containing common contaminants was used. The search was performed with tryptic cleavage specificity with 3 allowed miscleavages. Protein identification was under control of the false-discovery rate (FDR; <1 % FDR on protein and PSM level). In addition to MaxQuant default settings, the search was performed against the following variable modifications: Protein N-terminal acetylation, Gln to pyro-Glu formation (N-term. Gln) and oxidation (Met). Carbamidomethyl (Cys) was set as fixed modification. Further data analysis was performed using R scripts developed in-house. Missing LFQ intensities in the control samples were imputed with values close to the baseline. Data imputation was performed with values from a standard normal distribution with a mean of the 5 % quantile of the combined log_10_-transformed LFQ intensities and a standard deviation of 0.1. For the identification of significantly enriched proteins, the median log_2_ transformed protein ratios were calculated from four replicate experiments and boxplot outliers were identified in intensity bins of at least 300 proteins. Log_2_ transformed protein ratios of sample versus control with values outside a 1.5x (significance 1) or 3x (significance 2) interquartile range (IQR), respectively, were considered as significantly enriched. Volcano Plots were made based on the fold changes and p-values derived from the Bayesian test with robust correction on log_10_ scores using the limma package in R.

### Total proteomics

For proteome analysis, 25 μg of protein from each SILAC condition were pooled and proteins were reduced with 1 mM dithiothreitol. Samples were heated at 70°C for 10 min, alkylated by addition of 5.5 mM chloroacetamide, and loaded onto NuPAGE® Novex® 4-12 % Bis-Tris gels (Life Technologies). Proteins were separated by SDS–PAGE, stained using the Colloidal Blue Staining Kit (Life Technologies) and in-gel digested using sequencing grade modified trypsin (Sigma-Aldrich). Peptides were extracted from gel and desalted on reversed-phase C18 StageTips (Rappsilber et al. 2007). Peptides were analyzed on a quadrupole Orbitrap mass spectrometer (Exploris 480, Thermo Scientific) equipped with a UHPLC system (EASY-nLC 1200, Thermo Scientific) as described (Kelstrup et al. 2012; Bekker-Jensen et al. 2020). The mass spectrometer was operated in data-dependent mode, automatically switching between MS and MS2 acquisition. Survey full scan MS spectra (m/z 300–1,700) were acquired in the Orbitrap. The 15 most intense ions were sequentially isolated and fragmented by higher energy C-trap dissociation (HCD) (Olsen et al. 2007). An ion selection threshold of 5,000 was used. Peptides with unassigned charge states, as well as with charge states < +2, were excluded from fragmentation. Fragment spectra were acquired in the Orbitrap mass analyzer.

Raw data files were analyzed using MaxQuant (development version 1.5.2.8) (Cox and Mann 2008). Parent ion and MS2 spectra were searched against a database containing 96,817 human protein sequences obtained from the UniProtKB (February 2020 release) using the Andromeda search engine (Cox et al. 2011). Spectra were searched with a mass tolerance of 6 ppm in MS mode, 20 ppm in HCD MS2 mode, strict trypsin specificity and allowing up to two miscleavages. Cysteine carbamidomethylation was set as a fixed modification. N-terminal acetylation, oxidation, and N-ethylmaleimide (NEM) were set as variable modifications. The dataset was filtered based on posterior error probability (PEP) to arrive at a FDR < 1 % estimated using a target-decoy approach (Elias and Gygi 2010). Processed data from MaxQuant were analyzed in RStudio (version 4.1). Proteins or peptides flagged as “reverse”, “only identified by site” or “potential contaminant” were excluded from downstream analysis. Only proteins identified by no less than two peptides and at least one unique peptide were used and considered for downstream analysis. Statistical significance was assessed using LIMMA (Smyth 2004).

### Expression and purification of USP7, USP11-TCEAL1, TCEAL1

The genes for *H. sapiens* USP7 and USP11 were subcloned from AddGene vectors 46751 and 22566 into vector 438B (Gradia et al. 2017) via ligation independent cloning. The final constructs contain an N-terminal 6x His tag followed by a tobacco etch virus protease (TEV) cleavage site. *H. sapiens* TCEAL1 was cloned into vector 438A via ligation independent cloning (Gradia et al. 2017) and combined with USP11 to generate a co-expression vector. All proteins were expressed using baculovirus as previously described (Vos et al. 2016). All cells were grown in ESF-921 media (Expression Systems). Briefly, bacmids were isolated from DH10EMBacY cells (Geneva Biotech) and were transfected into Sf9 cells. Viruses were amplified in Sf21 cells to generate V1 virus. V1 virus was used to infect Hi5 cells for protein expression. 100-600 ml of Hi5 cells were infected with 50-300 µl of V1 virus and grown for 48-72 h after infection at 27°C. Cells were collected by centrifugation (238 x g), resuspended in lysis buffer containing 300 mM NaCl, 20 mM Na•HEPES pH 7.4, 10 % (v/v) glycerol, 30 mM imidazole pH 8.0 and 1 mM DTT, and snap frozen in liquid nitrogen.

All purification steps were performed at 4 °C unless otherwise noted. Frozen cells were thawed, lysed by sonication, and lysates were cleared by centrifugation. Clarified lysates were applied to Nickel-NTA resin (Takara) and incubated for 20 min with gentle rocking. Beads were washed 3 times with lysis buffer, 1 time with high salt buffer (lysis buffer with 1 M NaCl), followed by a final wash in lysis buffer. Protein was eluted with nickel elution buffer (Lysis buffer with 500 mM imidazole). Purified protein was quantified using the estimated extinction coefficient and absorbance at 280 nm. A Superose 6 3.2/300 column (GE) was equilibrated in 100 mM NaCl, 20 mM Tris-HCl pH 7.9, 4 % glycerol, and 1 mM DTT. 10 µM of USP7 or the USP11-TCEAL1 was applied to the column at a flow rate of 0.04 ml/min. Peak fractions were collected, separated by SDS-PAGE, and stained with Coomassie blue. USP7-USP11-TCEAL1 were mixed (11 µM final concentration) in a final buffer containing 100 mM NaCl, 20 mM Tris-HCl pH 7.9, 4 % glycerol, and 1 mM DTT and incubated at 30 °C for 30 min in a volume of 100 µl prior to application on the Superose 6 column. Peak fractions were collected, separated by SDS-PAGE, and stained with Coomassie blue.

### In vivo pulldown assays of recombinant USP11 constructs

Expression constructs coding for the USP11 catalytic domain (residues aa 298-963) and the N-terminal dUSP-Ubl1-catalytic domains (residues aa 70-963) were amplified from cDNA by PCR and cloned between the 3C site and the FLAG tag of pBADM-22-FLAG, an in-house vector carrying an N-terminal thioredixin-6xHis-tag and a C-terminal FLAG-tag under control of an Arabinose promotor by the SLIC-method (Li and Elledge 2007). These constructs were then used for the deletion of the catalytic domain insertion (residues aa 489-778) by plasmid linearization and subsequent recircularization by PCR using a single fusion-primer to bridge the resulting gap by a five residue linker AlaSerThrSerLys between residues 488 and 779 resulting in two further constructs coding for the core USP-domain (residues 298-488-ASTSK-779-963) and dUSP-Ubl1-core-USP-domain (70-488-ASTSK-779-963). All proteins were expressed in NEB 10-beta cells (New England Biolabs). Cells transformed with the corresponding expression vector were grown in LB medium supplemented with 100 µg/ml Ampicillin at 37 °C to an OD600 of 0.8. The temperature was then reduced to 18 °C and protein expression was induced by the addition of L-Arabinose to a final concentration of 50 µM. After an additional 16 h of shaking at 5 x g, the cells were harvested and resuspended in lysis buffer (50 mM HEPES pH 8.0, 300 mM NaCl, 1 mM TCEP, supplemented with Roche EDTA-free cOmplete protease inhibitor cocktail, DNaseI and Lysozyme). Cells were lysed using a cell disruptor and the lysate was cleared by centrifugation for 30 min at 35,000 x g The supernatant was applied to a gravity flow Ni-IDA column (Macherey & Nagel). The column was washed with 10 column volumes wash buffer (50 mM HEPES pH 8.0, 1 M NaCl) and proteins were eluted with lysis buffer supplemented with 400 mM Imidazole. The trx-His-tag was cleaved by the addition of 3C-protease during dialysis of the eluate against lysis buffer supplemented with 0.1 % (V/V) beta-mercaptoethanol overnight. Dialysates were then concentrated using spin-concentrators (10 kDa MWCO) and injected on a size exclusion column (Superdex 200 16/600, GE healthcare), pre-equilibriated with storage-buffer (20 mM HEPES pH 8.0, 200 mM NaCl, 1 mM TCEP). Fractions containing the desired protein were concentrated using spin-concentrators (10 kDa MWCO), aliquoted and flash-frozen in liquid nitrogen prior to storage at -80 °C until further use. SH-EP cells were lysed by HEPES lysis buffer (20 mM HEPES pH 7.9, 150 mM NaCl, 0.2% v/v NP-40, 0.5 mM EDTA, 10 % v/v Glycerol, 2 mM MgCl_2_) and 20 µg recombinant purified USP11 protein per ml cell lysate was added. Samples were incubated overnight at 4 °C on a rotating wheel. 20 µl/IP FLAG-M2 magnetic beads (Sigma-Aldrich) were washed with BSA-PBS (5 mg/ml) and added to the pulldown samples. As a control reference 1 % input per IP was kept. After 6 h at 4 °C circulating on a rotating wheel the beads were washed with HEPES buffer. Samples were eluted and subjected to western blotting as previously described.

### High-throughput sequencing

ChIP and ChIP-seq/ChIP-Rx was performed as described previously (Roeschert et al. 2021). For each ChIP-seq or ChIP-Rx experiment, 5x10^7^ cells per immunoprecipitation condition were fixed for 5 min at room temperature with formaldehyde (final concentration 1 %). The fixation reaction was quenched by adding 125 mM Glycine and incubating for 5 min at RT on an orbital shaker. After washing with PBS, each condition was collected in ice-cold PBS containing protease and phosphatase inhibitors (Sigma-Aldrich) and pelleted by centrifugation. All lysis buffer hereafter contained protease and phosphatase inhibitors. The cell pellets were resuspended in PIPES lysis buffer (5 mM PIPES pH 8.0, 85 mM KCl, 0.5 % v/v NP-40) and spiked with murine NIH-3T3 cells (ratio 1:10). After incubating for 20 min on ice the nuclei were pelleted by centrifuging at 252 x g for 5 min at 4 °C. The nuclear pellets were then resuspended in RIPA lysis buffer (50 mM HEPES pH 7.9, 140 mM NaCl, 1 mM EDTA, 0.1 % v/v SDS, 1 % v/v Triton X-100, 0.1 % w/v sodium deoxycholate) for 10 min at 4 °C. Chromatin was fragmented with a Covaris M220 Focused Ultrasonicator (Peak Power = 75.0, Cycles/Burst = 200, Duty Factor = 10.0, Duration = 3000 s per ml cell lysate, < 3*10^7^ cells/ml). Chromatin fragment size of 150-200 bp was checked by agarose gel electrophoresis. 1 % of the lysate was kept for input reference. A total volume of 30 µl (for ChIP) or 100 µl (for ChIP-Rx) per IP of 1:1 A/G Dynabeads mix (Thermo Fisher Scientific) was washed with 5 mg/ml BSA-PBS and incubated with 3 µg (for ChIP) or 15 µg (for ChIP-Rx) antibody against RNAPII (A-10), pSer2 RNAPII, TCEA1 or HA overnight at 4 °C on a rotating wheel. Chromatin samples were cleared by centrifugation at 21,952 x g for 20 min at 4 °C, and then incubated with the antibody/beads mix for 6 h at 4 °C on a rotating wheel. Then, the samples were washed three times each with washing buffer I (20 mM Tris-HCl 8.1, 150 mM NaCl, 2 mM EDTA, 1 % v/v Triton X-100, 0.1 % v/v SDS), washing buffer II (20 mM Tris-HCl pH 8.1, 500 mM NaCl, 2 mM EDTA, 1 % v/v Triton X-100, 0.1 % v/v SDS), washing buffer III (10 mM Tris pH 8.1, 250 mM LiCl, 1 mM EDTA, 1 % v/v NP-40, 1 % w/v sodium deoxycholate) and two times with TE buffer. The elution of the chromatin was done twice by incubating the beads with elution buffer (100 mM NaHCO_3_, 1 % v/v SDS in TE buffer) for 15 min at RT on a rotating wheel. Afterwards, input and immunoprecipitation samples were de-crosslinked first by digesting with RNase A for 1 h at 37 °C and incubation overnight at 65 °C with shaking, followed by digesting with proteinase K for 2 h at 45 °C with shaking. The isolation of the DNA was performed by phenol-chloroform extraction, purified with ethanol precipitation and finally quantified with the Quant-iT PicoGreen dsDNA assay (Thermo Fisher Scientific). Manual ChIP samples were subjected to RT-qPCR (see Table S2 for primer sequences). DNA libraries of ChIP-Rx samples were prepared by using the NEBNext Ultra II DNA Library Prep Kit for Illumina Sequencing (New England Biolabs).

RNA sequencing was performed by extracting RNA with RNeasy Mini Kit (Qiagen) according to the manufacturer’s instructions. On-column DNase I digestion was performed followed by mRNA isolation with the NEBNext Poly (A) mRNA Magnetic Isolation Kit (NEB). Library preparation was done with the Ultra II Directional RNA Library Prep for Illumina following the manufacturer’s manual. Libraries were size selected using SPRIselect Beads (Beckman Coulter) after amplification with 9 PCR cycles. Library quantification and size determination was performed with the Fragment Analyzer (Agilent) using the NGS Fragment High Sensitivity Analysis Kit (1-6,000 bp; Agilent). Libraries were subjected to cluster generation and base calling for 100 cycles paired end on the Illumina NextSeq 2000 platform.

### Immunofluorescence

Following treatment and incubation in 96-well plates (Perkin Elmer/Revvity) cells were washed and fixed with PFA (4 %) for 10 min at RT. Cells were then permeabilized with 0.3 % Triton X-100 for 10 min at RT and blocked with BSA-PBS (5 %) for 30 min at RT. Next, the cells were incubated with primary antibodies in blocking buffer overnight at 4 °C. After washing, cells were incubated with appropriate fluorophore-conjugated secondary antibodies for 1 h at RT and subsequently nuclei were counterstained with Hoechst33342 (2.5 µg/ml) for 10 min at RT. Image acquisition was done by using the Operetta CLS High-Content Analysis System with 40x magnification (PerkinElmer/Revvity) and images were processed using the Harmony High Content Imaging and Analysis Software. P values were calculated using an unpaired t-test comparing the mean fluorescence intensity of measured wells.

### In situ Proximity Ligation Assay

1,500-2,000 SH-EP cells expressing the doxycycline-inducible shRNA of interest were seeded per well in a 384 well format (PerkinElmer/Revvity). After doxycycline (1 mg/ml, 48 h) cells were fixed with 4 % PFA (Sigma-Aldrich) diluted in PBS for 30 min, washed with PBS and permeabilized with 0.3 % Triton-X-100 diluted in PBS for 20 min. After blocking (5 % BSA in PBS) for 1 h cells were incubated with primary antibodies in blocking solution overnight at 4 °C. After washing with PBS, PLUS (anti-rabbit) and MINUS (anti-mouse) probes (Sigma-Aldrich Duolink) were conjugated to primary antibodies for 1 h at 37 °C. The cells were then washed with Wash Buffer ‘A’ (Sigma-Aldrich Duolink) and the ligation reaction was carried out for 30 min at 37 °C, followed by washing and *in situ* PCR amplification for 2 h at 37 °C. Before washing with Wash Buffer ‘B’ cells were counterstained with Hoechst33342 (Thermo Fisher Scientific). Image acquisition was done using the Operetta CLS High-Content Analysis System with 40x magnification (PerkinElmer/Revvity) and images were processed using the Harmony High Content Imaging and Analysis Software. P values were calculated using an unpaired t-test comparing the mean signal per cell of measured wells.

### Structure prediction using AlphaFold

For 3D structure modelling of USP11 (UniProt: P51784) and TCEAL1 (Uniprot: Q15170) the AlphaFold Protein Structure Database (DeepMind, EMBL-EBI) was used. Further, to highlight specificity of complexes in detail the COSMIC^2^ web platform was used. Here, fasta files from UniProt entries were taken to run the AlphaFold-Multimer tool (v: 1.0) to predict 3D protein-protein complexes (Jumper et al. 2021; Varadi et al. 2022). For the multimer computations the default pretrained AlphaFold2 algorithm was used. The lowest predicted local distance difference test (pLDDT) scores to indicate prediction accuracy was used in Supplemental Figure 2b. Finally, the highest ranked AlphaFold Multimer prediction of five structures of the input set, sampled from five different model checkpoints, was evaluated using PyMOL (v: 2.5.5).

### Bioinformatics

Before high-throughput sequencing, the quality, quantity and size of the PCR-amplified DNA fragments of the prepared libraries were determined with a Fragment Analyzer (Thermo Fisher Scientific). All sequencing libraries were subjected to Ilumina NextSeq 500 sequencing according to the manufacturer’s instructions. After base calling with Illumina’s FASTQ Generation software (v1.0.0, NextSeq 500 Sequencing), high quality PF-clusters were selected for further analysis and sequencing quality was ascertained using FastQC software (v0.11.09; available online at: http://www.bioinformatics.babraham.ac.uk/projects/fastqc/). ChIP-Rx samples were mapped separately to the human hg19 and to the murine mm10 genome using Bowtie2 (v2.3.5.1 (Langmead and Salzberg 2012)) using the preset parameter “very-sensitive-local”. Further, ChIP samples were normalized to the number of mapped reads in the smallest samples. Human ChIP-seq samples were either normalized relative to the spiked-in mouse reads (ChIP-Rx), or to the same number of human reads (read-normalized samples). The normalized bam files were sorted using SAMtools v1.9 (Danecek et al. 2021) and converted to bedgraphs using bedtools genomecov (v2.30.0 (Quinlan and Hall 2010)). Integrated Genome Browser (Nicol et al. 2009) was used to visualize these density files. Metagene and density plots were generated with ngs.plot.r v2.63-7 (Shen et al. 2014) or DeepTools v3.5.1 (Ramirez et al. 2016) using a bin size of 10 bp. P-values were calculated by comparing the mean of the reads count via unpaired *t*-test for the specified genomic position showed in the average density plots. Plots labeled with “all genes” refer to the 57,773 genes annotated for hg19/GRCh37.p13 by Ensembl v75 (Feb 2014). Pausing index were calculated by the ratio between RNAPII density in the promoter (TSS -30 to +300 bp) and in the gene body region (TSS +300 bp to TES). Only genes were included with a median RNAPII read density >0.01 over the gene body and/or the TSS. RNA-seq samples were mapped to human genome hg19 with STAR version 2.7.10a (Dobin et al. 2013). Differential gene expression was calculated with edgeR v3.28.0 using biological triplicates for each condition.

### mRNA gene expression signature analysis

Genes downregulated upon TCEAL1 depletion were Pearson correlated with the mesenchymal signature scores (Groningen et al 2017) in the SEQC neuroblastoma cohort with 498 neuroblastoma samples (RNAseq). Signature scores were analyzed in the R2 Genomics Analysis and Visualization Platform (https://r2.amc.nl) using the unidirectional (all mesenchymal signature genes are ‘elevated’) rank-version signature calculation. In short, for every sample in the 498 neuroblastoma data set, genes were sorted on the level of expression. Then within this ranked gene order, every signature gene is looked up, while recording the ratio position. The average ratio of the complete signature genes forms the signature score for that sample. This procedure is repeated for every sample. These signature values are used as a metagene to correlate against all the downregulated genes.

## Acknowledgements

This work was supported by grants from the Mildred Scheel Junior Research Center Program (to G.B.), the European Research Council (SENATR ERC #101096948 to M.E.), the German Cancer Aid (#70114538 to M.E), the German Research Foundation (EI 222/21-1 and INST 93/1023-1-FUGG to M.E. and project-ID 464588647 - SFB 1551 to P.B.), the Alex’s Lemonade Foundation Crazy 8 Initiative (to G.B., S.M.V. and M.E.), Berlin Institute of Health and Stiftung Charité (Ro.V.), Fight Kids Cancer Funding Program (Ro.V. and J.K.). S.M.V. is a Freeman Hrabowski Scholar of the Howard Hughes Medical Institute. The authors thank Barbara Bauer, André Kutschke and Tobias Roth for technical support.

## Author Contributions

Conceptualization, G.B. and M.E.; Methodology, G.B., P.B., S.M.V. and M.E.; Investigation, M.D., S.H., G.C., F.C., F.S., C.P.A., C.S.-V. and G.B.; Formal Analysis, J.K., Ra.V. and P.G.; Visualization, M.D., P.G. and G.B.; Writing – Original Draft, G.B. and M.E.; Supervision, P.B., C.K., Ro.V., S.M.V., M.E. and G.B.; Funding Acquisition, G.B., S.M.V., P.B., J.K., Ro.V. and M.E.

## Data Accessibility

Sequencing datasets are uploaded at the Gene Expression Omnibus. We will provide the accession number GSE and the reviewer access token as soon as we have received feedback. The proteomics data have been deposited to the ProteomeXchange Consortium via PRIDE (Vizcaino et al. 2013). Reviewers can access the datasets by logging in to the PRIDE website using the following account details: Project accession: PXD052970 Token: B6waj25NMRj3.

**Supplemental Table 1:**

Excel File summarizing results of mass spectrometry.

---

**Supplemental Figure 1:**
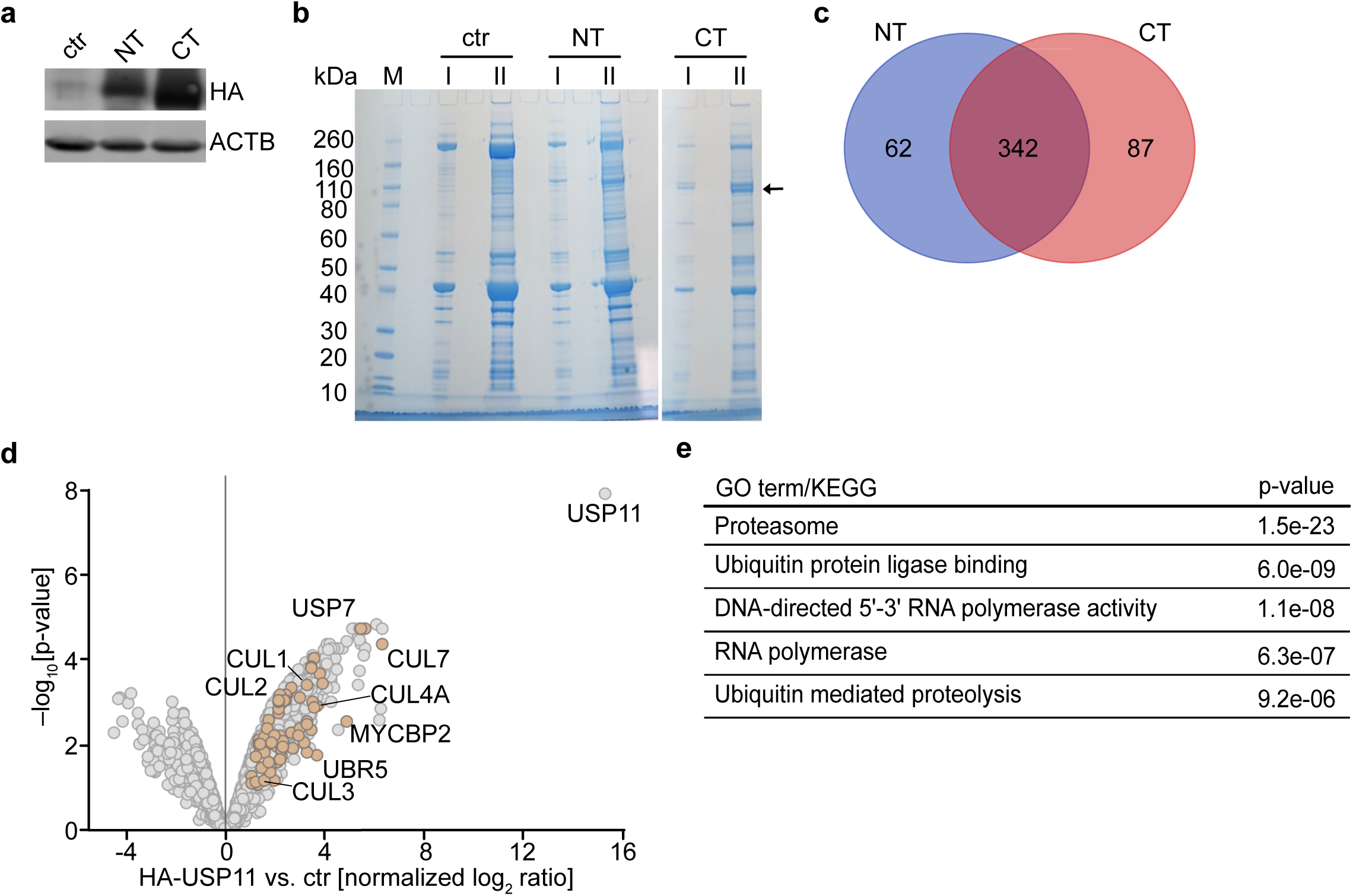
Identification of the USP11 interactome. **a.** Immunoblot of SH-EP cells expressing N- or C-terminally HA-tagged ectopic USP11 (NT or CT, respectively). ACTB was used as a loading control (n=2). **b.** Coomassie stained SDS gels showing 15 % (I) and 85 % (II) of samples used for MS-based USP11 interactome analysis. The black arrow indicates the size of the HA-tagged USP11 protein. M marks the molecular weight marker lane. **c.** Venn diagram showing overlap of USP11 interactors co-immunoprecipitated with either NT- or CT-tagged HA-USP11 in SH-EP cells. **d.** Volcano plot of the consensus USP11 interactome with proteins involved in the ubiquitin system marked in brown. The x-axis displays the log_2_ ratio of proteins in HA-USP11 IPs relative to control IPs. The y-axis shows the -log_10_ of p-value for each protein. **e.** GO-terms significantly enriched in the USP11 interactome relative to all proteins.

**Supplemental Figure 2:**
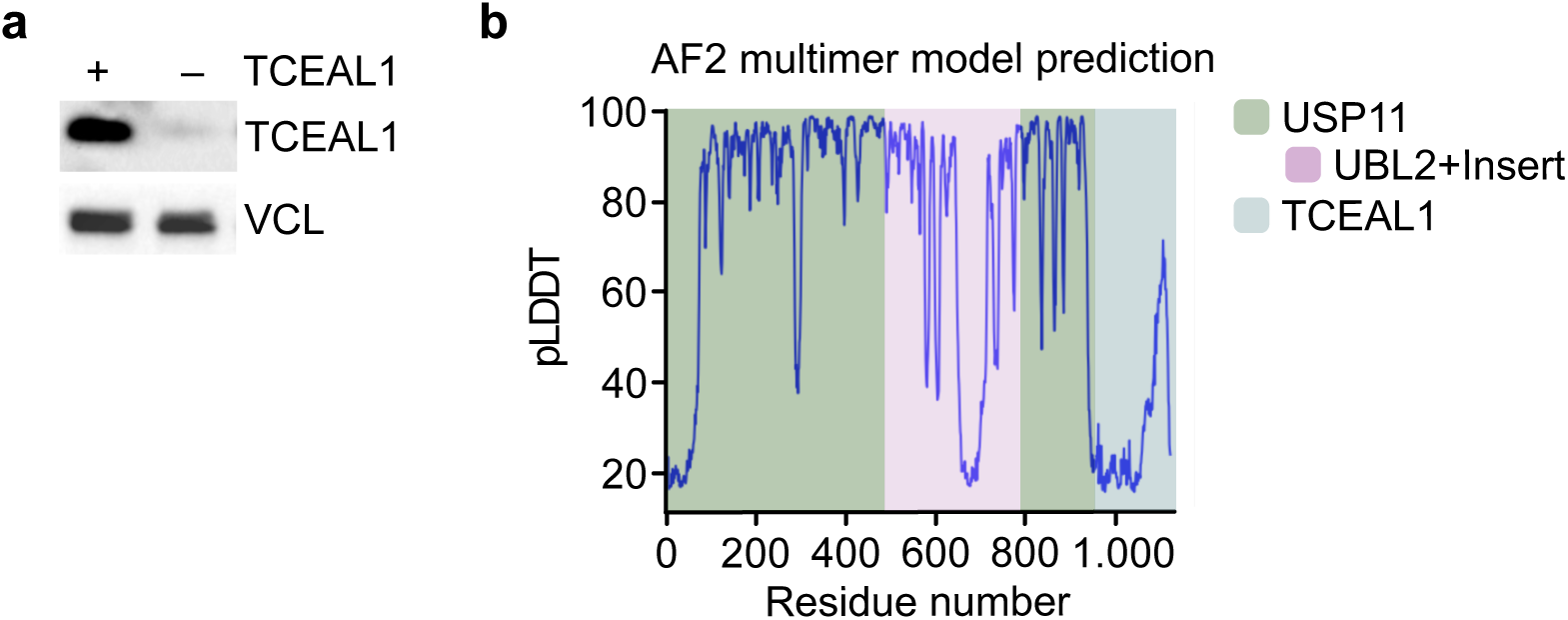
UBL2+Insert domain in USP11 is needed for complex formation with TCEAL1. **a.** Immunoblot showing levels of TCEAL1 in SH-EP cells expressing doxycycline-inducible shRNAs targeting *TCEAL1*. Where indicated cells were treated with doxycycline (1 µg/ml, 48 h). VCL was used as loading control (n=3). **b.** Model prediction of USP11 structure (green/purple) and TCEAL1 (blue) multimer by AlphaFold2 algorithm. The y-axis displays the per-residue confidence metric predicted local distance difference test (pLDDT) score (0-100). The x-axis displays the residue number of the multimer given by the order of FASTA sequence input.

**Supplemental Figure 3:**
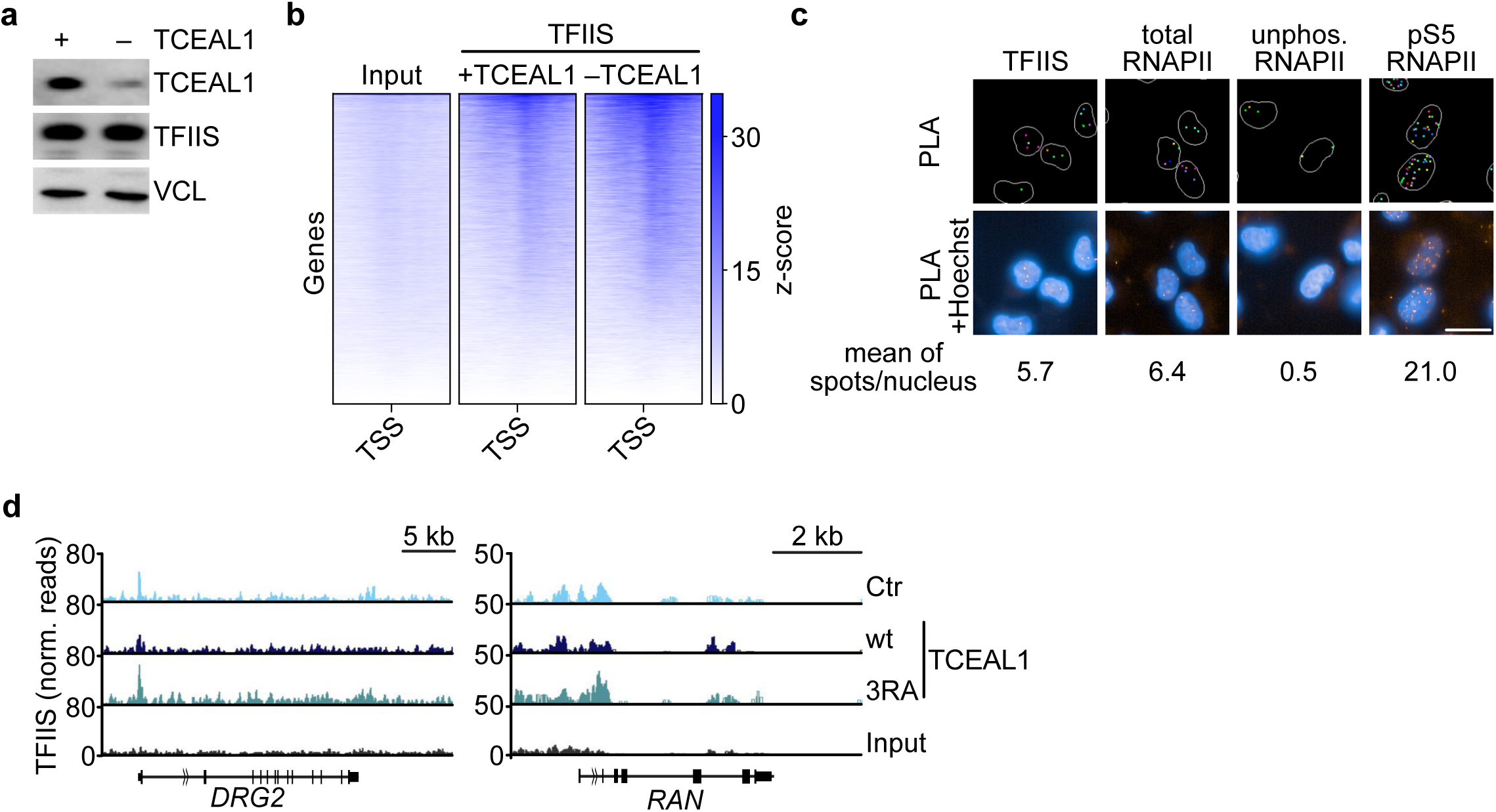
TFIIS accumulates at highly expressed genes in TCEAL1-depleted cells. **a.** Immunoblots showing levels of TCEAL1 and TFIIS in SH-EP cells expressing doxycycline-inducible shRNAs targeting *TCEAL1* (1 µg/ml, 48 h). VCL was used as loading control (n=3). **b.** Heatmap of TFIIS chromatin binding centered to the transcription start side (TSS) upon TCEAL1 depletion (N=16,285 genes are shown). **c.** Representative images of spot detection in single antibody controls of PLAs shown in Figure 4f. The numbers indicate the mean of PLA spots per nucleus in these cells. Scale bar indicates 20 µm (n=3). **d.** Genome browser tracks of indicated loci showing chromatin association of TFIIS upon expression of TCEAL1^wt^ or TCEAL1^3RA^. Cells without ectopic TCEAL1 were used as control (Ctr) (n=2).

**Supplemental Figure 4:**
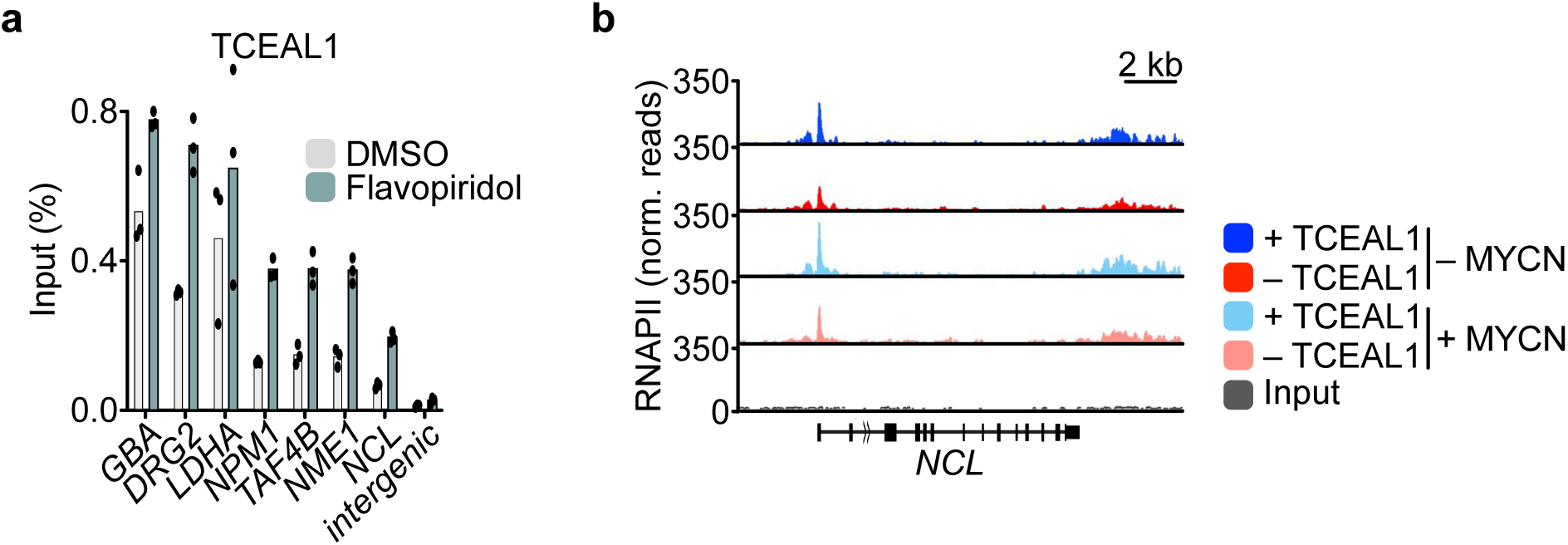
TCEAL1 stabilizes elongating RNAPII. **a.** HA-TCEAL1 ChIP in SH-EP cells treated with flavopiridol or DMSO at indicated promoters. Shown is the mean of technical triplicates of one representative experiment (n=3). **b.** Genome browser tracks of indicated locus showing chromatin association of RNAPII in SH-EP-MYCN-ER cells in control cells (blue) or upon doxycycline-mediated depletion (red) of TCEAL1 without (top) or with (bottom) MYCN induction.

**Supplemental Figure 5:**
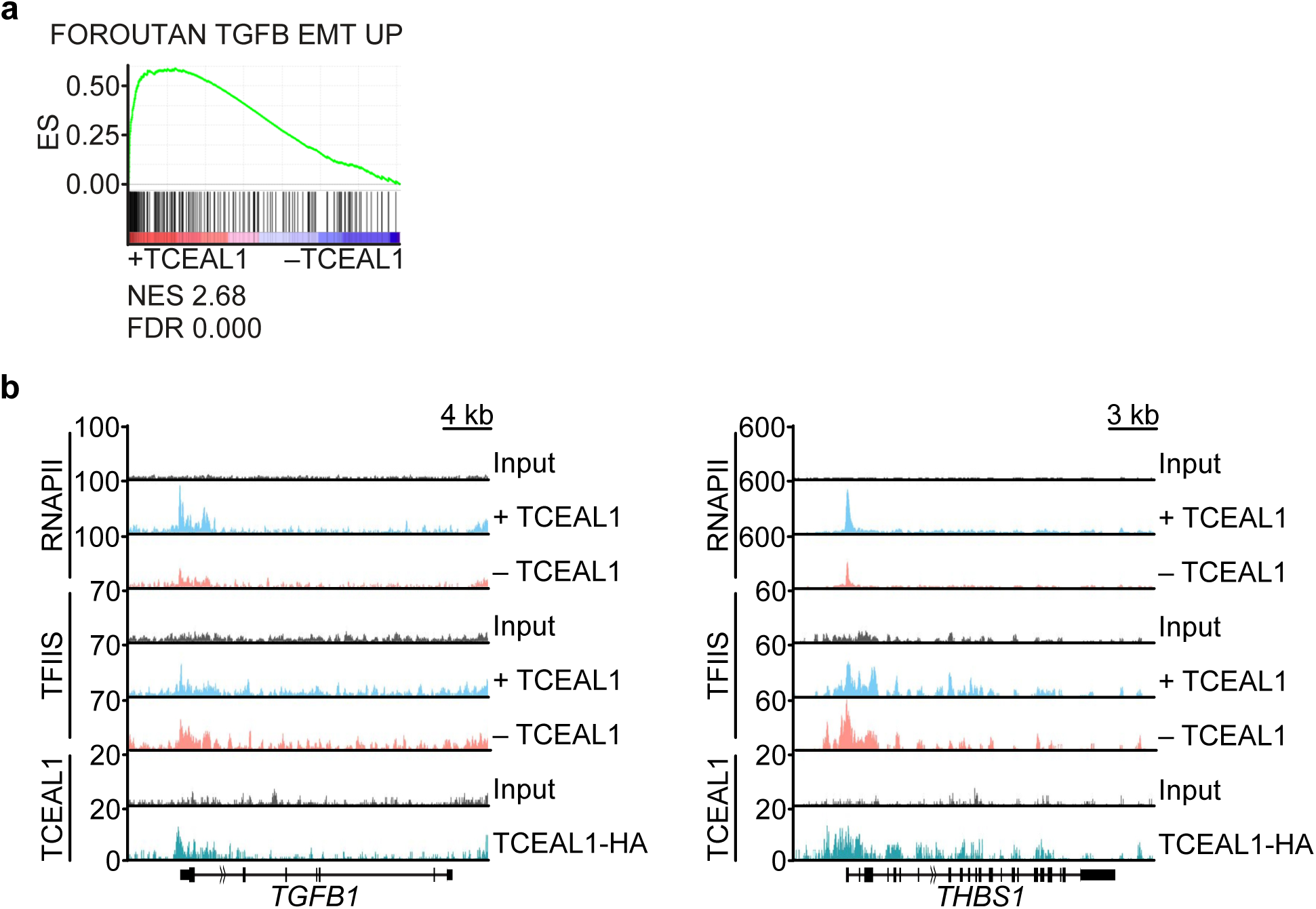
**a.** GSEA plot of the indicated gene set upon TCEAL1 depletion in SH-EP cells. NES, normalized enrichment score; FDR, false discovery rate. **b.** Genome browser tracks shown as normalized reads of ChIP-seq for the indicated proteins at the *THBS1* (encoding thrombospondin 1) and the *TGFB1* (TGF beta1) gene loci. Input belongs to the different sequencings.

